# Centennial recovery of recent human-disturbed forests

**DOI:** 10.1101/2024.07.21.604432

**Authors:** Asun Rodríguez-Uña, Verónica Cruz-Alonso, José A. López-López, David Moreno-Mateos

## Abstract

International commitments are challenging countries to restore their degraded lands, particularly forests. These commitments require global assessments of recovery timescales and trajectories of different forest attributes to inform restoration strategies. We use a meta-chronosequence approach including 125 forest chronosequences to reconstruct past (*c*. 300 years), and model future recovery trajectories of forests recovering from agriculture and logging impacts. We found recovering forests significantly differed from undisturbed ones after 150 years and projected that difference to remain for up to 218 (38-745) or 494 (92-2,039) years for ecosystem attributes like nitrogen stocks or species similarity, respectively. These conservative estimates, however, do not capture the complexity of forest ecosystems. A centennial recovery of forests requires strategic, unprecedented planning to deliver a restored world.

## Main Text

Global commitments accumulate aims to restore world’s degraded ecosystems. Most of these commitments are now backboned by three major initiatives: the UN Decade on Ecosystem Restoration that delineated major lines of action (*1*), the outcomes from COP 15 of the Convention on Biological Diversity that goes a step further creating the largest restoration fund ever planned, accumulating 600 billion dollars (*2*), and the European Nature Restoration Law, the first continent-wide legally binding regulation where restoration is explicitly required by all EU countries (*3*). This rapidly increasing demand to reverse biodiversity loss through restoration has, however, critical unresolved questions related to the promise of a restored world (*4, 5*). Ecosystem recovery from anthropogenic disturbance is a long-term process involving the reassembly of an unknown number of interactions that generate the functions required for the ecosystem to persist through time and respond to further disturbances. As environmental conditions change, the ability of ecosystems to respond to disturbance can be altered, and may globally diminish in the case of some forests (*6*). Given the timescale and the complexity of this process, it remains unclear whether we can deliver a restored world within the next few generations.

Over the last two decades, global restoration assessments have consistently showed restored ecosystems hosting lower biodiversity and reduced rates of biogeochemical cycling than undisturbed ones (*7, 8*). Specific assessment in rivers (*9*), wetlands (*10*), marine ecosystems (*11*), or forests (*12*) follow similar patterns. These assessments have, however, provided inconsistent results on the factors that affect recovery performance. Specifically, for example, the effects of the restoration approach taken or the effect of the initial degrading activity. It is unclear if this limited recovery of restored ecosystems would fade as recovery progresses, and if it does so, when levels of biodiversity and function similar to those found today in less disturbed areas will be attained. Thus, attaining similar levels of complexity (in terms of species interactions) involving previously non-present in the restored ecosystem, or non-dominant species, and the functions linked to those interactions is critical (*4, 13*). Unless these challenges are addressed, restoration risks adding to the current biotic homogenization or simplification (less species and interactions more broadly distributed) (*14*).

Despite the potential relevance of forest restoration to decelerate biodiversity loss and accumulate carbon (*15, 16*), most restored forests to date do not recover to levels of biodiversity and functioning similar to those existing in reference less disturbed sites (*7, 12, 17*). A comprehensive mostly neotropical forest recovery study reported incomplete recovery of plant biomass and species composition after 120 years (*17*). There have been contrasting results about the additional benefits provided by forest restoration actions compared to natural regeneration (*12, 18*–*20*). Previous studies have suggested longer times to recovery after disturbances of greater intensity like large-scale clear-cutting, than after small-scale disturbances like shifting cultivation (*12, 18, 21*). It remains, however, unclear *i)* the effect of the restoration approach and the disturbance type on the recovery trajectory; and *ii)* the extent at which recovery varies as a function of the metric used to track ecosystem change.

One of the factors that may compromise forest recovery and restoration success is our limited understanding of the recovery patterns, and mechanisms, at timescales relevant for ecosystems commonly dominated by species with centennial lifespans. The lack of studies monitoring recovery using direct observations at those timescales can be partially addressed by the use of space-for-time substitutions (i.e., chronosequences) to reconstruct recovery trajectories (*22, 23*). We use a unique meta-chronosequence approach (*12, 24*–*26*) that includes complete recovery trajectories of forests worldwide under contrasting conditions (i.e. restoration strategy, disturbance type, climate) that may critically shape recovery trajectories. This allowed us to build *c*. 300-years composite recovery trajectories of forests recovering through passive or active restoration approaches, after two of the most widespread impacts on forests, agriculture and logging. We included six recovery metrics related to biodiversity (i.e., organism abundance, species diversity, species similarity) and biogeochemical functions (i.e., carbon cycling, nitrogen stock and phosphorus stock) (*27*). This study allowed *i)* characterizing the recovery trajectory and calculating the recovery completeness reached by forests at ecologically meaningful timescales as affected by the recovery metric used; *ii)* estimating the time to recovery for each metric; and *iii)* understanding the effect of human-related (i.e., disturbance type, restoration strategy) and environmental (i.e., aridity) factors on the recovery process. Our results provide information about the real extent of forest degradation, the subsequent timescales at which forest recovery operates and some of its drivers that can be considered in global, regional and national restoration strategies and environmental agendas.

### Meta-analysis predictors

We collected data from 16,873 plots from 125 chronosequences of recovering forest ecosystems in 110 published primary studies, including a total study area >183,500 km^2^ (see materials and methods; supplementary text; figs. S1 and S2; tables S1 and S2) (*27*). From these chronosequences, we extracted 641 recovery trajectories of quantitative measures of ecosystem attributes along time, related to six recovery metrics (organism abundance, species diversity, species similarity, carbon cycling, nitrogen stock, and phosphorus stock), two restoration strategies (passive and active), three disturbance types [agriculture (including abandoned croplands and pastures), logging and mining], and a climatic metric (i.e., aridity index) (*27*). The trajectories related to organism abundance mainly contained biomass and density measurements. Species diversity included measurements of species richness and diversity indexes. Species similarity trajectories contained information about species composition along the recovery trajectory, which were used to calculate pairwise compositional similarity at specific recovery times compared to a reference value. We used the Morisita-Horn similarity index, which accounts for species relative abundance (*28*). Abundance, diversity and composition trajectories included five life forms: plants (including trajectories of woody plants, non-woody plants, and all plants combined), invertebrates, microorganisms, fungi and birds. Carbon cycling included pools and fluxes in soil, plants, litter and microorganisms, whereas nitrogen and phosphorus stocks included pools in soil, plants and litter (table S2). From each trajectory, we collected all available recovery measures and compared them with a reference value. This analysis resulted in 3,400 quantitative comparisons, computed as response ratios that divide values of the same metric in recovering and reference forests (*27*). Reference values were selected from old-growth forests (66%) or, if unavailable, secondary forests >100 years old (34%) with similar environmental conditions.

### Forest recovery completeness and time to recovery

While biodiversity and biogeochemical functions of forests recovering globally from agriculture and logging reached different levels of recovery after 150 to 300 years, it would take more than 200 to 500 years to reach levels similar to those existing in reference, less disturbed sites, according to our models (*27*) (Figs. 1 and 2 and fig. S3). The initial condition of the forest, determined by the intercept of the models, had a strong positive effect on recovery completeness (fig. S4 and tables S3 and S4). This suggests that the intensity of the impacts exerted in a forest can determine the length of its recovery process. Recovery trajectories followed, however, different trajectories. Abundance of organisms, species similarity and nitrogen stocks followed a logarithmic function, whereas carbon cycling and species diversity followed a square root function (*27*) (Fig. 1A and fig. S3 and table S5). This suggests that the recovery of abundance of organisms, species similarity and nitrogen stocks may be faster, but not necessarily more complete, than the recovery of carbon cycling and species diversity during the first 150 years after the end of the disturbance regardless of the initial condition of the forest. Phosphorus stocks followed a linear recovery trajectory suggesting less adverse initial conditions, but slower recovery, than carbon cycling and nitrogen stocks (Fig. 1A and fig. S3 and tables S4 and S6). Our trajectory analysis shows that recovery responds to many more factors than the ones we have used in this study, and suggests that substantially more metrics would be needed to understand the mechanisms of forest recovery and design effective tools to accelerate the process in restoration actions.

**Fig. 1.**
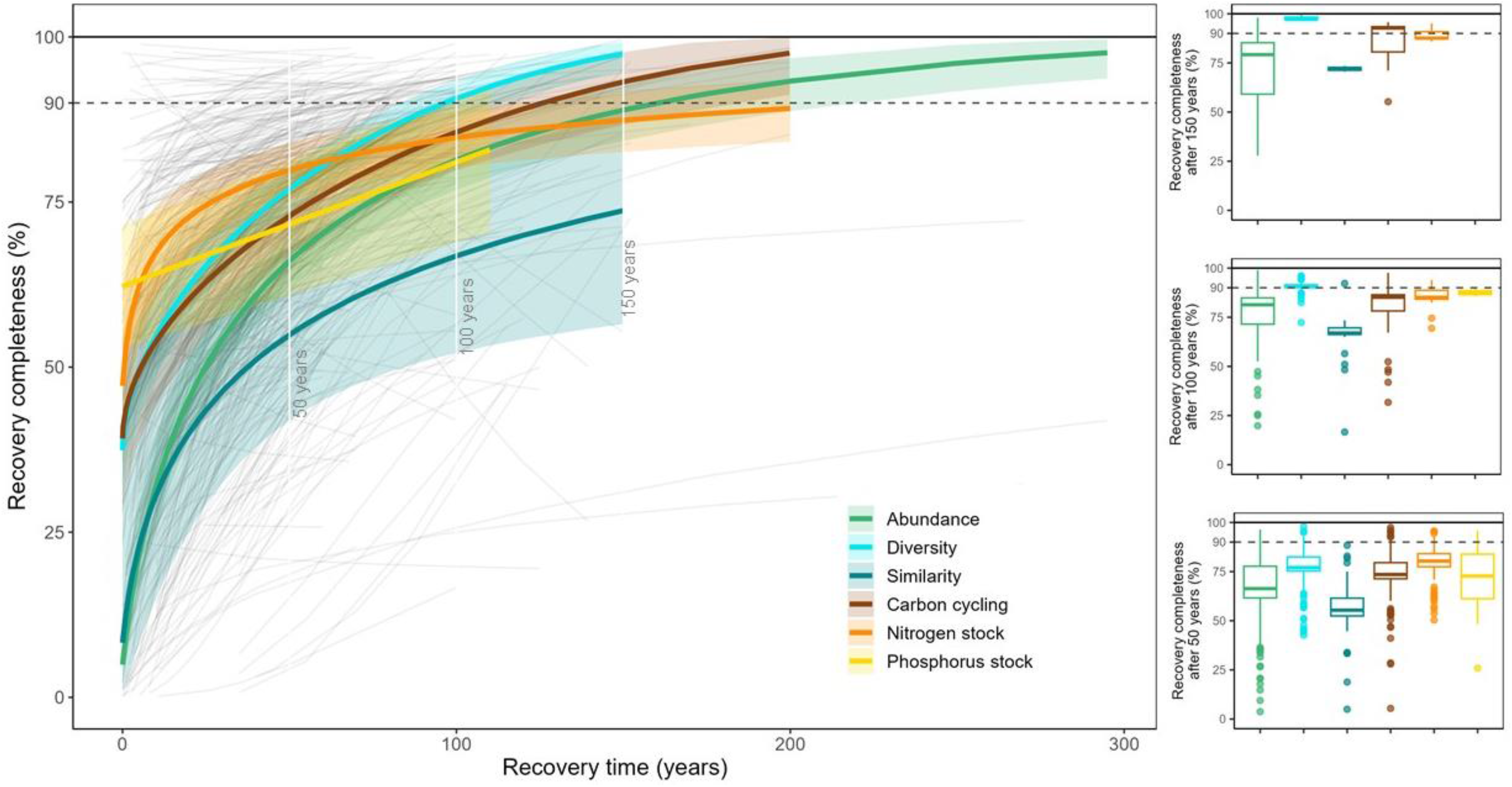
Recovery completeness of forest biodiversity and functions. (**A**) Recovery trajectory of forest biodiversity (i.e., organism abundance, species diversity and species similarity) and functions (i.e., cycling of carbon, nitrogen stock and phosphorus stock). Thick colored lines of each recovery metric correspond to a consensus trajectory in which different intercepts and slopes were fitted for each individual trajectory (thin grey lines). Shadowed areas include the 95% confidence intervals of the fixed effects. Recovery completeness was computed with log-response ratios. The dashed and solid horizontal lines represent the recovery of 90% and 100% of the reference goal value, respectively. (**B**) Boxplots of the predicted recovery completeness of each metric after 50 years (lower panel), 100 years (middle panel) and 150 years (upper panel), based on the models used for Fig. 1A. The dashed and solid horizontal lines represent the recovery of 90% and 100% of the reference goal value, respectively.

**Fig. 2.**
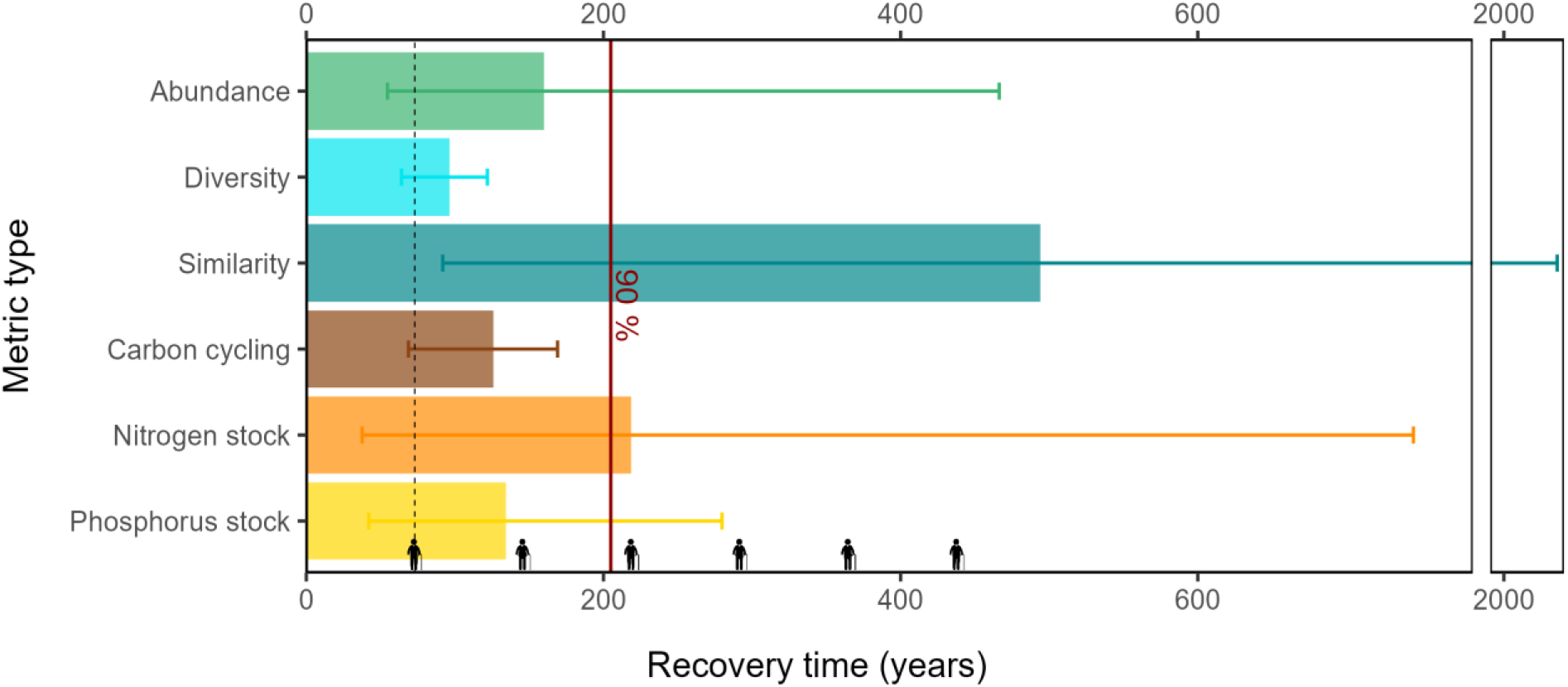
Time to forest recovery. Predicted time to reach > 90% recovery of forest biodiversity (i.e., organism abundance, species diversity and species similarity) and functions (i.e., cycling of carbon, nitrogen stock and phosphorus stock). Error bars indicate 0.10-0.90 quantiles. The red solid line represents the predicted average time to reach > 90% recovery. The dotted black line indicates the global human life expectancy (i.e., 73 years) and the elderly person icons indicate subsequent human generations.

The predictions of our models fitted for the effect of recovery time on the response ratio [mean (95% confidence interval)] after one, two and three times the global human life expectancy (i.e., 73 years) showed that the recovery of organism abundance may be incomplete [94.5% (90.0-97.7%)] after three human generations (*27*) (219 years; Fig. 1A and fig. S3 and table S7). After one human generation, the estimated recovery ranged from 61.3% (48.0-74.6%) for species similarity to 84.5% (81.3-87.4%) for species diversity. Following those projections, we estimated that it would take 160 (55-466) [median (0.10-0.90 quantile ranges)] years for organism abundance, 97 (64-122) years for species diversity, 494 (92-2039) years for species similarity, 126 (69-169) years for carbon cycling, 218 (38-745) years for nitrogen stocks and 134 (42-283) years for phosphorus stocks to recover to 90% of the reference old-growth values (*27*) (Fig. 2 and table S8).

These projections based on available data bring forward previous estimations of forest recovery time (*17, 24, 29, 30*) by several centuries. Specific metrics, e.g. carbon cycling, align however with previous estimations related to the recovery of biomass (*17*). Our projected recovery times support the hypothesis that old-growth forests are irreplaceable in terms of the biodiversity and functions they host (*30, 31*) and adds that recovery will likely take no less than 500 years only considering our simple recovery metrics. We suggest expanding the complexity of the metrics used to assess forest recovery, which could include metrics related to the structure and stability of species interaction networks, its interactions with the phylogenetic structure of the community and their functional responses (*4, 32*). These metrics require less likely combinations of elements (e.g., phenological synchronicity, trait match) that could lead to longer, more accurate recovery times (*4, 33*). Finally, the fact that 34% of the selected studies included a secondary forest of at least 100 years as a reference may also be artificially contributing to shorter, more conservative recovery times compared to studies that used actual old-growth forests.

### Forest recovery predictors

Our models on recovery completeness 50 years after the disturbance showed higher recovery when forests were recovering from being meadows formerly grazed than from being logged, cultivated or cultivated and grazed combined (Fig. 3 and tables S9 and S10; up-to-50-year data allowed for robust statistical analysis). Also, the models by metric showed that after 50 years *i)* the abundance of forest organisms and their diversity recovered 1.00-1.06 and 1.00-1.03 (95% confidence interval) times more from grazing than from logging impacts, respectively; *ii)* species similarity recovery between recovering and reference forests was 1.05-1.31, 1.00-1.32 and 1.01-1.31 times higher after grazing than after logging, cultivation, or combined cultivation and grazing impacts, respectively; *iii)* carbon cycling measurements recovered 1.00-1.06 and 1.01-1.05 times more from grazing and cultivation than from logging impacts, respectively; and *iv)* nitrogen cycling measurements recovered 0.98-1.04 times more from grazing than from logging impacts (Fig. 3 and tables S11 – S17). Even after 100 years, forests had higher (1.03-1.34) recovery from cultivation than from logging for their carbon cycling measurements (fig. S5A; tables S14). These results indicate that the impact of logging activities on forests may last for longer than the impacts of cultivation and grazing. This unexpected pattern may be caused by agricultural legacy effects leaving high soil nitrogen and phosphorus pools (*34, 35*) that would help plants and other soil organisms overcome common nitrogen or phosphorus limited conditions (*36, 37*). These new conditions would support plant establishment and growth (*38*) and linked carbon accumulation rates and processes controlled by biomass production and decomposition (*39*). An unreported accumulation of clear-cut forest practices, where large amounts of nutrients and organic carbon are removed, could alternatively help explain the logging effects. However, we could not test this hypothesis with the available data in the studies included in this assessment.

**Fig. 3.**
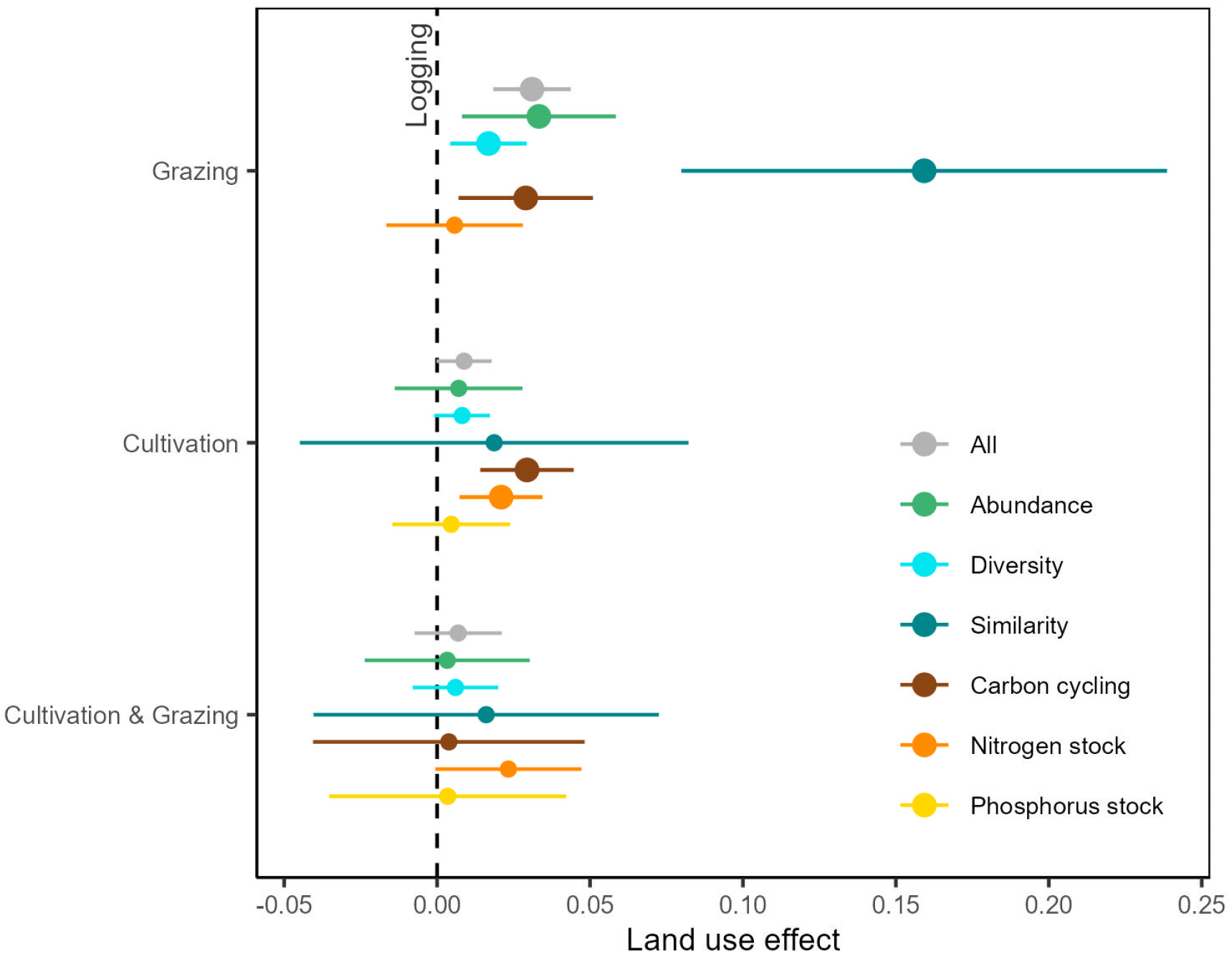
Effect of previous disturbance on forest recovery. Effect of the previous disturbance category on the recovery completeness of forest biodiversity (i.e., organism abundance, species diversity and Morisita-Horn species similarity) and functions (i.e., cycling of carbon, nitrogen stock and phosphorus stock) after 50 years since recovery started. Recovery completeness was computed with log-response ratios. Error lines represent 95% confidence intervals of the estimated effect. Those error lines not overlapping the zero dashed line (shown with bigger dots) correspond to statistically significant effects. This means a higher recovery completeness after cultivation, grazing or combined cultivation and grazing than after logging (the reference category in the model) when there is a positive effect.

We did not detect significant differences between actively or passively restored forests through time in any of the metrics used in this study (fig. S5B and fig. S6A; tables S11 – S16). This result expands previous studies comparing passive and active restoration approaches (*12, 19*) suggesting that the lack of effects of actively restored forests compared with abandoned lands persists even after century of recovery. By accounting for the effect of the initial condition of the forest (*27*), we could overcome the common site selection bias linking highly degraded areas to more active restoration strategies (*40*). We did not observe differences either among life forms (e.g. woody vs. non-woody plants; fig. S5D and fig. S6C; tables S9, S11–S13). This suggests that the nature of the metrics (e.g. abundance, diversity) rather than the organism measured may have a stronger response to the recovery effects. This supports the need of a diversity of ecosystem, rather than organism, focused metrics to increase the accuracy of recovery assessments. We also found that recovery overall was particularly challenged in arid environments even after 100 years of recovery, where drier forests had lower abundance of organisms than wetter forests (fig. S6C; tables S11 – S16).

Large scale restoration efforts are currently developed in arid regions to reverse land degradation as the need to increase plant cover in those regions is critical to regulate water flows, avoid erosion or improve soil functionality (e.g., *41, 42*). Our results suggest that recovery may fall short in arid lands compared to wetter regions and highlight the need to devote additional resources to the restoration of arid lands to reach similar chances of recovery to those existing in wetter regions.

## Applications

Our suggested centennial recovery of forests affected by agriculture and logging highlights the need to plan forest restoration strategies and environmental regulations at the relevant timescales and adapted to the initial conditions (i.e. post-agricultural or post-logging land). This information is particularly critical at a moment where large scale strategies, particularly the UN Decade on Ecosystem Restoration and regional implementation strategies (e.g., AFR100 in Africa, Initiative 20×20 in Latin America and the Caribbean), are expecting to deliver restored forests on a scale of decades. A centennial recovery also highlights that old-growth forests, harboring unique specialists and functions, are irreplaceable in periods likely beyond a few centuries. We recommend caution in promoting restoration to reverse biodiversity and functional loss of forests in detriment of protection (e.g. in the context of biodiversity offsetting programs) as this could increase the amount of less-functional and diverse forest ecosystems. To better align our restoration promises with the reality of forest recovery, there is an opportunity to develop strategic planning at the timescales we have found relevant for forest recovery and to further restoration research. We suggest a research agenda focusing on the recovery of forest complexity at relevant timescales and the practical tools required to accelerate the recovery of that complexity.

## Acknowledgments

We are grateful to June Hidalgo for helping with the construction of the database and to Andrew Tanentzap and Serguei Saavedra for their comments.

## Funding

Environmental Fellowship Program (2016) of Tatiana Pérez de Guzmán el Bueno Foundation (ARU)

Spanish State Research Agency through María de Maeztu Excellence Unit accreditation 2018-2022 MDM-2017-0714 (DMM, ARU)

Real Colegio Complutense Postdoctoral Fellowship (VCA)

## Author contributions

Funding acquisition: DMM

Supervision: DMM

Conceptualization: DMM, ARU

Data curation: ARU

Methodology: ARU, VCA, DMM, JALL

Formal Analysis: VCA, ARU, JALL

Visualization: VCA

Writing – original draft: ARU, DMM, VCA

Writing – review & editing: ARU, VCA, DMM, JALL

### Competing interests

Authors declare that they have no competing interests.

### Data and materials availability

All data and code used in this study will uploaded to the Dryad digital data repository once the article is accepted for publication (link).

## Supplementary Materials

Materials and Methods

Supplementary Text

Figs. S1 to S8

Tables S1 to S17

References (*43–186*)

## Notes

### Competing Interest Statement

The authors have declared no competing interest.

