## Supplementary materials for "Centennial recovery of recent human-disturbed forests"

**Contents:**

Materials and Methods

Supplementary Text

Figs. S1 to S8

Tables S1 to S17

References (43–186)

### Materials and Methods

Study selection

We conducted a literature search in January 2020 in the scientific database ISI Web of Science using the search chain "(chronoseq* and forest* and (recover* or restor* or rehab* or regen*) not fire)". We excluded fire disturbance because all studies in a preliminary search corresponded to natural fires. The search produced 862 results, but 476 were rejected after a title and abstract screening (fig. S1 for PRISMA flow diagram (*43*)). From the remaining studies, we selected those that *i)* included forest recovery information for at least 50 years; *ii)* had at least two measurements of recovery in time to be compared with a pre-disturbance reference, an old-growth forest or a mature forest that have been recovering for at least 100 years (hereafter, reference forest); *iii)* reported time since recovery started; *iv)* were directly related to any anthropogenic disturbance (except fire); and *v)* included temporal trajectories of organism abundance, species diversity, species similarity, carbon cycling, nitrogen stock or phosphorus stock. We excluded less common recovery metrics, mainly related to biogeochemical functions such as soil pH, to reduce database heterogeneity. Our selection yielded 110 published primary studies including 125 chronosequences of recovering forest ecosystems (figs. S1 and S2 and table S1).

Database construction

From the selected chronosequences, we extracted 641 recovery trajectories, i.e., field-based quantitative measurements of ecosystem integrity repeated through time, reported in tables, figures and text of the selected studies. Each trajectory included at least two data-points, defined as the value of the ecosystem metric at different times since recovery started (hereafter, recovery time). Average values were considered for the data-points with the same recovery time (n = 72, in 21 studies). We used the free software Engauge Digitizer version 11.2 (*44*) to extract the data from the figures. To increase the homogeneity of the database, prior to data extraction, we carried out an initial training test by three people entering data. The test consisted of simultaneous data extraction of the same three studies and a posterior agreement on discrepancies. Afterwards, every two weeks the extraction team met to solve further discrepancies.

Our selection included trajectories whose recovery time ranged from 50 years to 295 years (84 ± 1 years, mean ± s.e.). The trajectories were related to agriculture (n = 370), logging (n = 243) or mining (n = 2) disturbances. Agricultural trajectories were subdivided in cultivation (n = 221), grazing (n = 90) and combined cultivation and grazing (n = 59, when forests were recovering in both abandoned croplands and pastures). For 26 trajectories, both agriculture and logging practices took place and no information was provided about if they occurred simultaneously or not. The trajectories related to organism abundance (n = 209) contained measurements of: *i)* biomass, density or cover of trees, shrubs or herbs, *ii)* height, diameter or basal area of trees, and *iii)* biomass or density of bacteria, fungi or invertebrates. The trajectories related to species diversity (n = 160) included measurements of species richness or diversity indexes, e.g., Shannon and Fisher's alpha. Species similarity trajectories (n = 42) contained information about species composition (i.e., species lists and their occurrence, total or relative abundance or frequency) at different recovery times along the chronosequence, which were used to calculate pairwise compositional similarity between each data-point and the reference value. We used three common similarity metrics: Morisita-Horn and Bray-Curtis indices, which account for species relative abundance, and Jaccard index, which considers species occurrence (*45*). Abundance, diversity and similarity trajectories included six life forms: woody plants (n =

254), non-woody plants (n = 33), woody and non-woody plants combined (n = 17), invertebrates (n = 72), microorganisms (n = 18), fungi (n = 15) and birds (n = 2). The cycling of carbon included pools (n = 114) in soil (n = 82), plants (n = 17), litter (n = 17) and microorganisms (n = 1), and fluxes (n = 14), i.e., rates of decomposition, respiration or sequestration in soil (n = 9), plants (n = 1) or litter (n = 4). The trajectories related to nitrogen stocks (n = 77) and phosphorus stocks (n = 25) included pools in soil (n = 78), plants (n = 15) and litter (n = 9) (table S2).

For each trajectory, we also recorded latitude and longitude, area of the study site, number of data-points along the trajectory, number of reference sites, number of plots sampled per trajectory and their area, number of subplots (i.e., replicates within plots) and their area, and restoration strategy [passive (n = 567) *vs*. active (n = 74)]. We found data from 16,873 plots, whose size ranged from 0.01 to 50 ha. The sizes of the study areas (reported in 89% of the chronosequences), ranged from 0.13 ha to 24,716 $km^2$. We estimated a total accumulated study area >183,500 $km^2$ (table S2). With the coordinates of each chronosequence, we extracted mean annual temperature and precipitation from WorldClim 2.1 (http://worldclim.org). We calculated the Lang aridity index (*46*) by dividing the precipitation by the temperature. Further database construction details are described in the Supplementary Text.

<u>Effect size calculations</u>

We used response ratios (RRs) to estimate the recovery completeness, i.e., the effect sizes between reference and recovering systems (*47*, *48*). We computed the RR for each data-point along the trajectory as $\ln(\frac{X_{res}}{X_{ref}})$, where $X_{res}$ is the value of the ecosystem metric at a certain recovery time and $X_{ref}$ is the reference value of the same metric in the reference forest. The reference value was from a nearby old-growth forests for 64% of the trajectories or the same forest ecosystem in the pre-disturbance state for 2% of the trajectories. None of these options were available for 34% of the trajectories. In these cases, we used a secondary, long-term recovery stage by selecting the last available data-point beyond 100 years on the trajectory.

In 32% of the RR computed, recovering values exceeded reference values ($X_{res} > X_{ref}$). This was especially the case in early recovery stages for measurements of plant density, species richness and soil nitrogen concentration. Metrics from ecosystems recovering from disturbances may show higher variability than in pre-disturbance states (*49*, *50*), providing values of those metrics that go above reference values for extended periods of time (*51*, *52*). The fact that ecosystem performance values are above the reference do not necessarily involve better, nor worse, ecosystem performance, it involves a deviation from the reference. For this reason, we give those values the same weight than the ones below the reference and consider all as deviations from the reference. In these cases, we computed the inverted RR as $\ln(\frac{X_{ref}}{X_{res}})$ (*53*, *54*). In the case of similarity trajectories, the similarity indices calculated (i.e., Morisita-Horn, Bray-Curtis and Jaccard) already represent a response metric between reference and recovering systems. Yet, their values needed to be standardized to account for the different scales at which beta-diversity may occur in different studies. Therefore, for seven (23%) chronosequences with at least two reference data-points, we computed the RR as $\ln(\frac{Sim_{res}}{Sim_{ref}})$, where $Sim_{res}$ is the value of the similarity index at a certain recovery time and $Sim_{ref}$ is the average value of the similarity index of all reference data-points in that chronosequence. For the remaining 24 chronosequences without reference variability, we used the same value of $Sim_{ref}$ for all of them, equal to the average of the seven average reference similarities previously calculated. The overall average reference similarity of Morisita-Horn, Bray-Curtis and Jaccard indices was 0.63 ± 0.27%, 0.52 ±

0.14% and 0.36 ± 0.13% (mean ± SE), respectively. RRs cannot be calculated for data-points or references with zero values; thus, we excluded 67 (1.9%) comparisons where $X_{res} = 0$ (n = 41), $X_{ref} = 0$ (n = 15) or $Sim_{res} = 0$ (n = 11). In total, we accumulated 3,400 RRs, i.e., quantitative comparisons between reference and successional recovery stages.

Weighting

Effect sizes in meta-analyses are usually weighted by study precision, which is typically the inverse of the variance. However, as in most ecological meta-analyses (*47*, *55*, *56*), the data necessary to determine variance was absent in the majority (72%) of our trajectories. Alternatively, we estimated the precision as the product of the number of subplots and their area, assuming that a higher sampling effort would imply a higher precision (*57*). The sources of variation for the RR varied between biodiversity and biogeochemical functions, where within-study heterogeneity and among-study heterogeneity account for most of the variation, respectively. For abundance, diversity and similarity, the value of among-study heterogeneity ($\tau^2$) was zero, meaning that fixed- and random-effects weights were equivalent, and then the heterogeneity statistic $I^2$ index (*58*) also equaled zero. In these cases, we fitted fixed-effects models, with weights only accounting for within-study variability. On the contrary, $I^2$ values were 96%, 49% and 41% for the RR of carbon, nitrogen and phosphorus, respectively, suggesting the presence of heterogeneity for those outcomes. Furthermore, their random-effect meta-analytic model fitted the data better than the fixed-effect meta-analytic model in terms of the Akaike Information Criterion corrected for small samples (AICc) (*59*). Thus, we assumed random-effect meta-analytic models for biogeochemical functions, accounting for both between- and within-study variation. To improve model convergence, the magnitude of the weights was reduced by dividing them by the constant $e$ for carbon and by $e^2$ for abundance, diversity and similarity.

Statistical analysis

To estimate the trajectory of forest recovery along time, we fitted a separate linear mixed model (LMM) for the RR of each recovery metric. We included the recovery time as fixed factor and as a random slope, and the trajectory identity as random intercept, enabling a different slope and intercept for each trajectory. As the recovery process along time may result in a wide range of trajectories from linear to more saturating shapes (*60*), we consider three functions to include the recovery time variable: one linear and two decelerating trends [ln(recovery time + 1) and √recovery time]. We then selected among the three options the one that best fit to the data of each recovery metric according to the minimum AICc (Table S5). To fit the models, 20 RR values were removed from the abundance dataset, all belonging to the same chronosequence, with a disproportionally large plot size compared with the rest of the data. Two RR values with the lowest values of recovery completeness and clearly discordant with other RR values (i.e., -6.9 and -7.8) of the diversity dataset were also removed. The models for the recovery of similarity were fitted using the Morisita-Horn index, as the Pearson correlation test informed that it was correlated to Jaccard and Bray-Curtis indices (fig. S7). Resulting RR values ranged from -8.5 to 0. Their absolute values were square root transformed to meet the assumptions of general linear models, and then multiplied by -1 to facilitate interpretation.

Using the resulting LMMs, we predicted the RR after 73, 146 and 219 years of recovery [i.e., one, two and three times the global life expectancy in 2019 (*61*)]. We then predicted the time needed for forest ecosystems to recover to 90% of reference values for each trajectory and recovery metric and calculated the median by metric. Reaching 100% recovery might be theoretically unfeasible, given the decelerating nature of the functions that best explained recovery trajectories of most recovery metrics. It can also be ecologically unfeasible due to natural variability, the uniqueness of each ecosystem, and the effects of global changes that would prevent an exact match of the reference system (*62*).

Also using the resulting LMMs, we predicted the RR after 50 and 100 years of recovery for each metric and trajectory (1) to know if the recovery completeness is dependent on the metric and (2) to understand the main explanatory variables underlying the recovery process for each metric. We fitted liner models (LM) to analyse the difference in the RR after 50 years and after 100 years of recovery among recovery metrics. The models had the recovery metric and the intercepts of the LMMs for each trajectory as fixed factors. The latter was included to account for the effect of the initial state of degradation when recovery started (fig. S4). We then fitted a separate LM for the effect of each explanatory variable studied (i.e., aridity, disturbance category, restoration strategy or life form) on the RR predictions after 50 and 100 years of all recovery metrics together, and then for each recovery metric individually. In all the cases, the intercept of the LMMs for each trajectory was also included as fixed factor to account for the effect of the initial state of degradation when recovery started. For the models fitted for the disturbance category and the life form, we excluded the categories with <1% of the values (i.e., "mining" for disturbance and "bird" for life form) or those including data with mixing information from other categories (i.e., "agriculture and logging" for disturbance and "woody and non-woody" for life form).

Models were fitted using R 3.6.3 (*63*) package *lme4* (*64*). In all cases, we standardized continuous explanatory variables by subtracting the mean and dividing by the standard deviation (*65*). We evaluated the good-of-fitness of each model by both comparing the observed and the fitted values ($r^2$ calculated with *performance* package (*66*); fig. S8) and AICc was computed using *MuMIn* package (*67*). The *p*-values of the effects in LMMs were estimated with *lmerTest* package (*68*). We used *emmeans* package (*69*) to define the differences among metrics, disturbance categories and life forms when they were significant. Model predictions were estimated using *ggeffects* package (*70*) and linear model assumptions checked using *performance* (*66*). Data manipulation and plotting was facilitated by *readxl* (*71*), *googledrive* (*72*), *tidyverse* (*73*), *broom* (*74*), *patchwork* (*75*) and *sjPlot* (*76*) packages.

### Supplementary Text

Database construction

The extraction of some variables from the studies required additional considerations as follows.

*Plot area*. When trajectories were obtained by sampling all the trees inside a given a plot or all the trees with a diameter at breast height higher than a given dimension (n = 51, in 7 studies), we assigned a number of subplots equal to one. When soil cores were extracted and no information was reported about their dimensions (n = 29, in 9 studies), we assumed a diameter of 10 cm, as it was the most common measure in the rest of cores registered. When a hole was dug and the dimensions were not provided (n = 14, in 5 studies), we assigned the most common dimensions for the other holes registered, which were 30 x 30 cm. When transects were used and only the length was provided (n = 12, in 3 studies), we assumed that 1 m was sampled to the right and to the left of that line as it was the most common width used in the rest of the studies.

*Recovery time*. When the recovery time was reported as an interval, we considered the average value of that interval (n = 228, in 14 studies). When the recovery time was reported as an open interval with values above or below a specific number (n = 42, in 7 studies) we used that exact number.

*Disturbance type*. In chronosequences with a history of different types of disturbances (e.g., forest was first clear-cut and then cultivation occurred), we considered the last disturbance prior to the beginning of forest recovery.

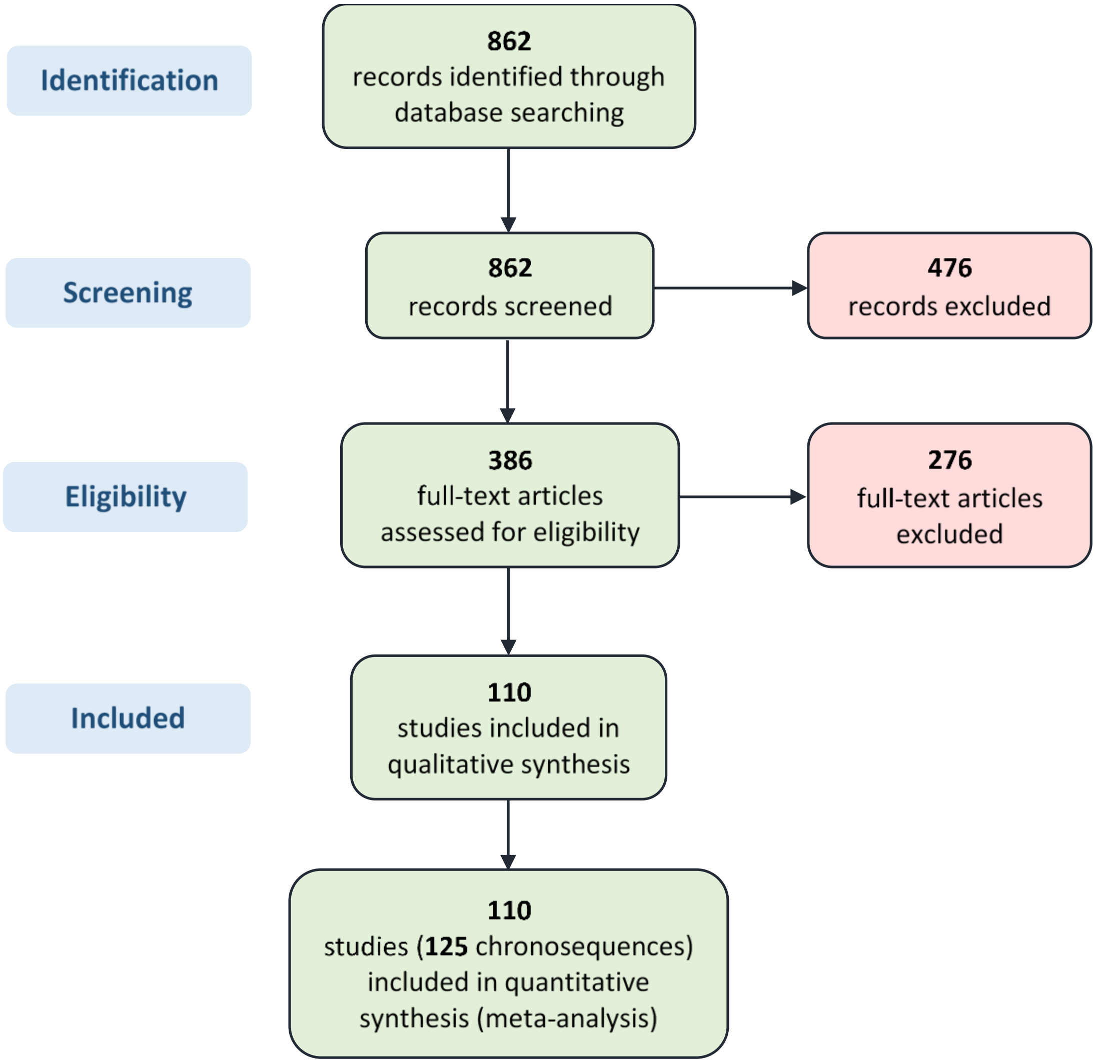


**Fig. S1.** Structure and template for flow chart from (*43*).

A

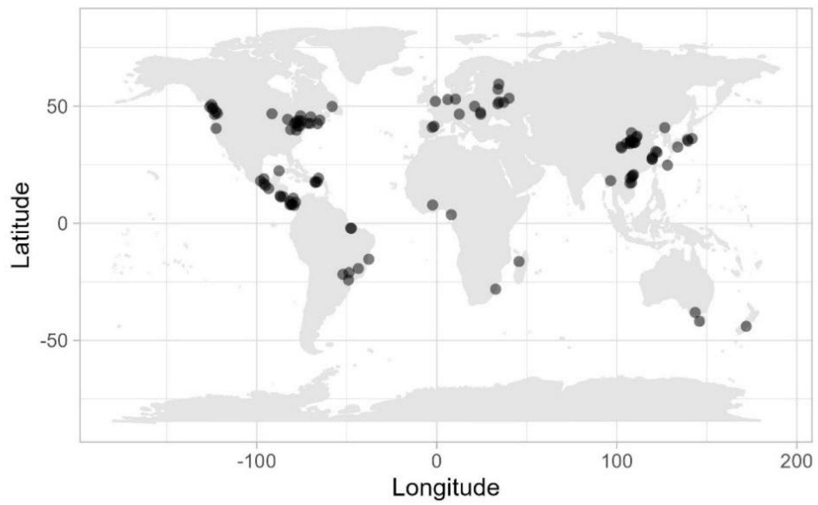


B

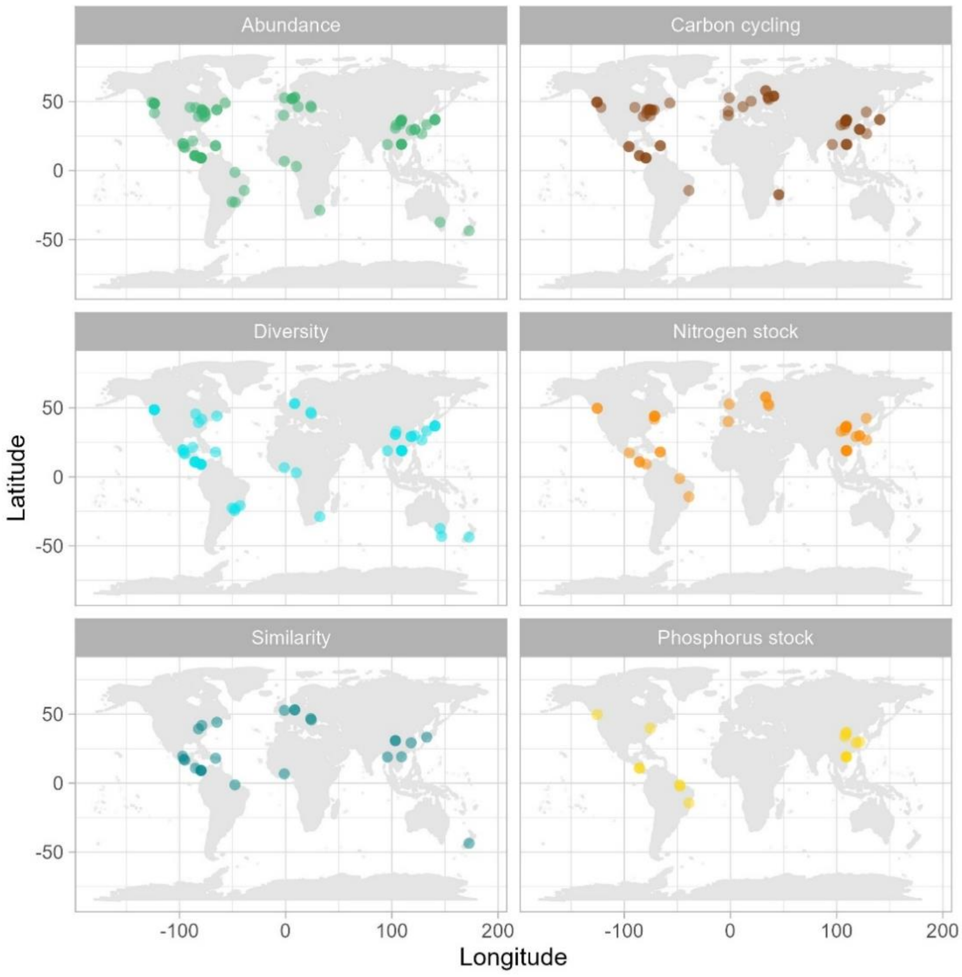


**Fig. S2.** Global distribution of (A) the 125 chronosequences studied and (B) the chronosequences depending on the recovery metrics (i.e., organism abundance, species diversity and Morisita-Horn species similarity, cycling of carbon, nitrogen stock and phosphorus stock).

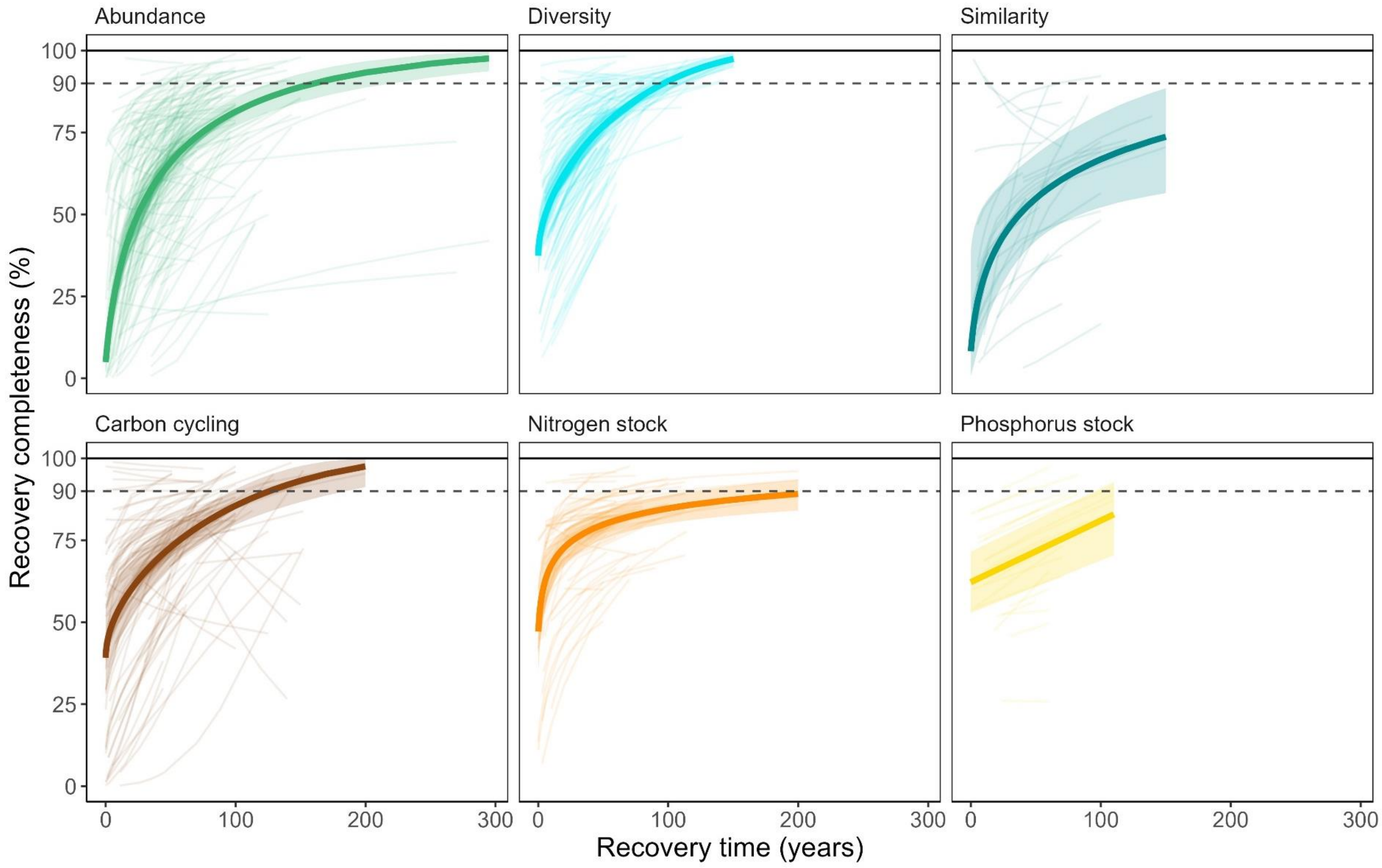


**Fig. S3.** Recovery trajectories of forest biodiversity (i.e., organism abundance, species diversity and Morisita-Horn species similarity) and functions (i.e., cycling of carbon, nitrogen stock and phosphorus stock). Thick line of each recovery metric corresponds to a model in which different intercepts and slopes were fitted for each trajectory (thin lines). Shadowed areas include the 95% confidence intervals of the fixed effects. The dashed and solid horizontal lines represent the recovery of 90% and 100% of the reference goal value, respectively.

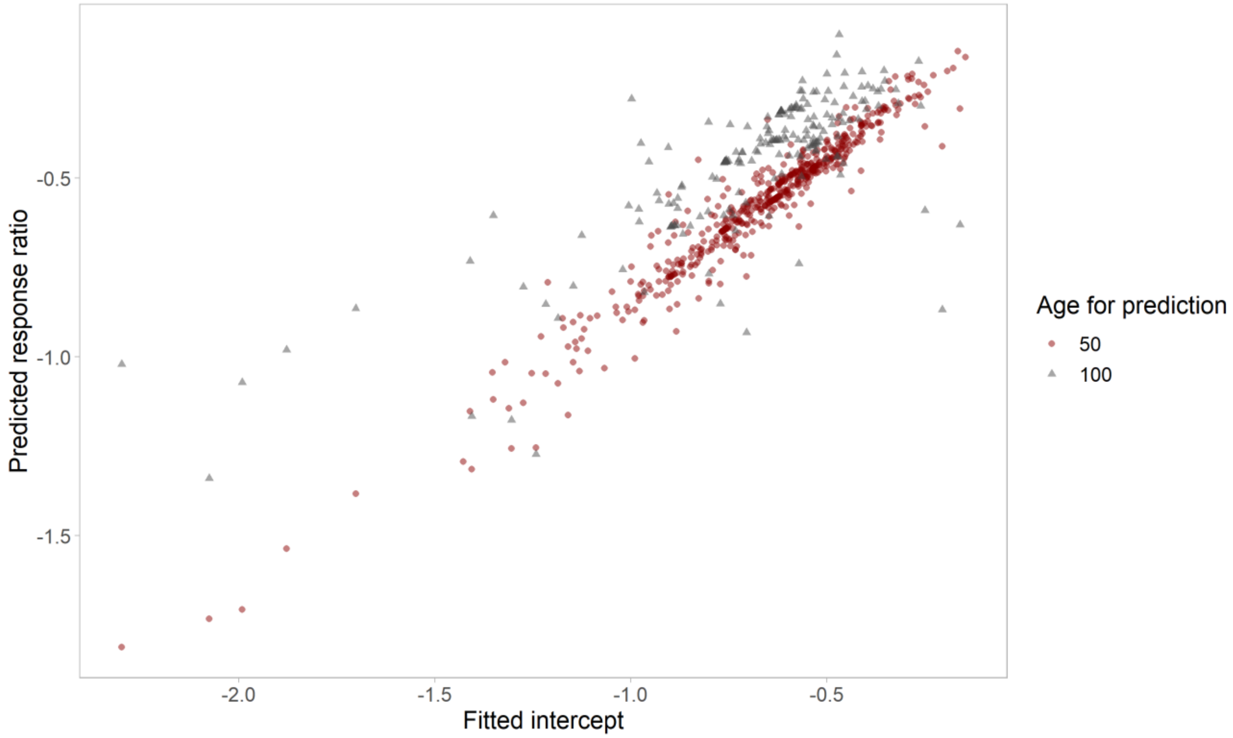


**Fig. S4.** Relationship between the predicted response ratio after 50 and 100 years of recovery and the intercept of each trajectory for all recovery metrics. The intercepts were obtained from the mixed models fitted for the effect of recovery time on the response ratio for each recovery metric.

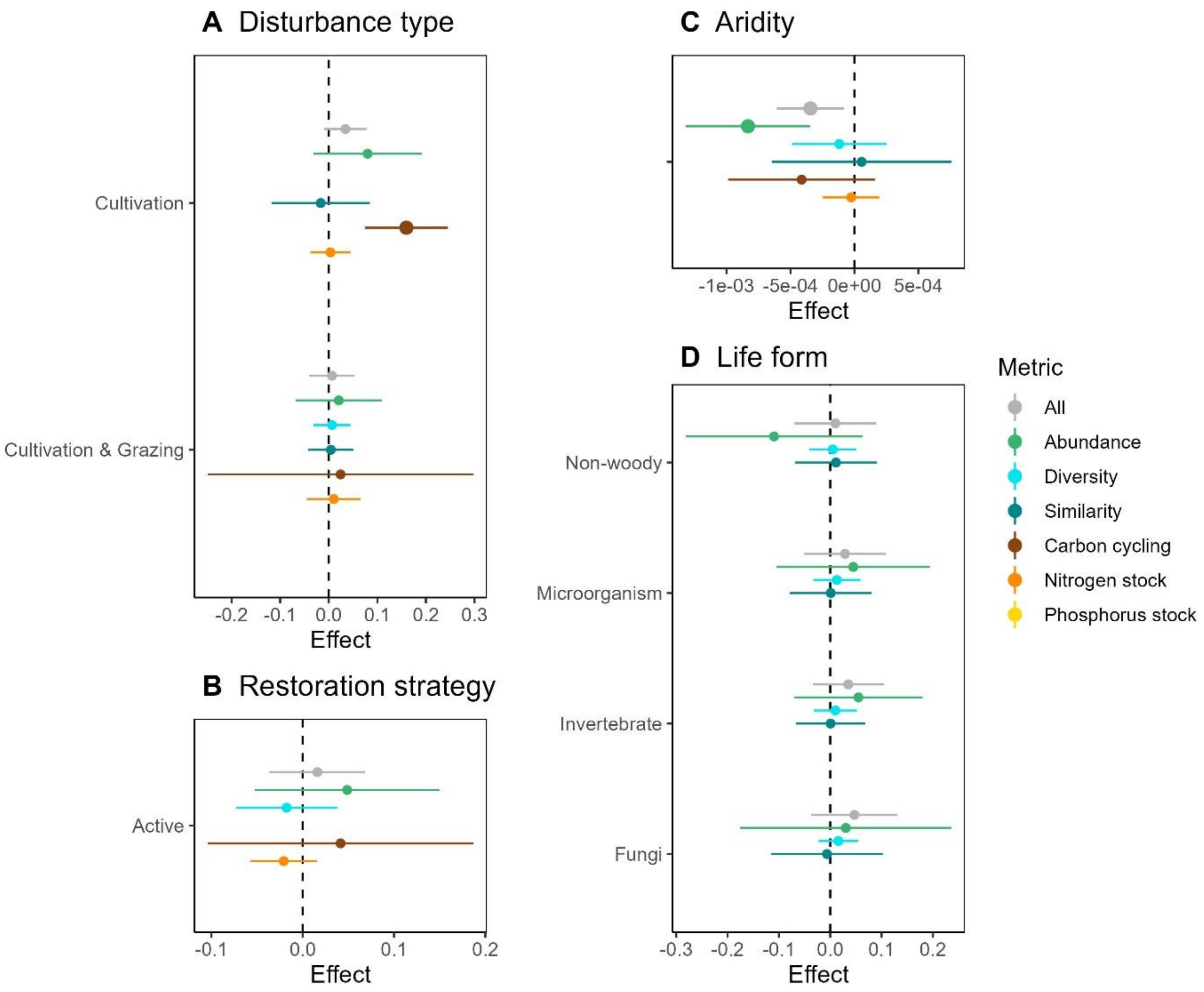


**Fig. S5**. Effect of the disturbance category (A), restoration strategy (B), aridity (C) and life form (D) on the recovery completeness of forest biodiversity (i.e., organism abundance, species diversity and Morisita-Horn species similarity) and functions (i.e., cycling of carbon, nitrogen stock and phosphorus stock), after 100 years since recovery started. Recovery completeness was computed with log response ratios. Error lines represent 95% confidence intervals of the estimated effect. Those error lines not overlapping the zero dashed line (shown with bigger dots) correspond to statistically significant effects. The reference categories for categorical variables were "logging" (for disturbance category), "passive" (for restoration strategy) and "woody" (for life form). Having a positive effect means a higher recovery completeness after cultivation or combined cultivation and grazing than after logging, with active restoration than with passive, and when the specific organism group is measured instead of woody plants.

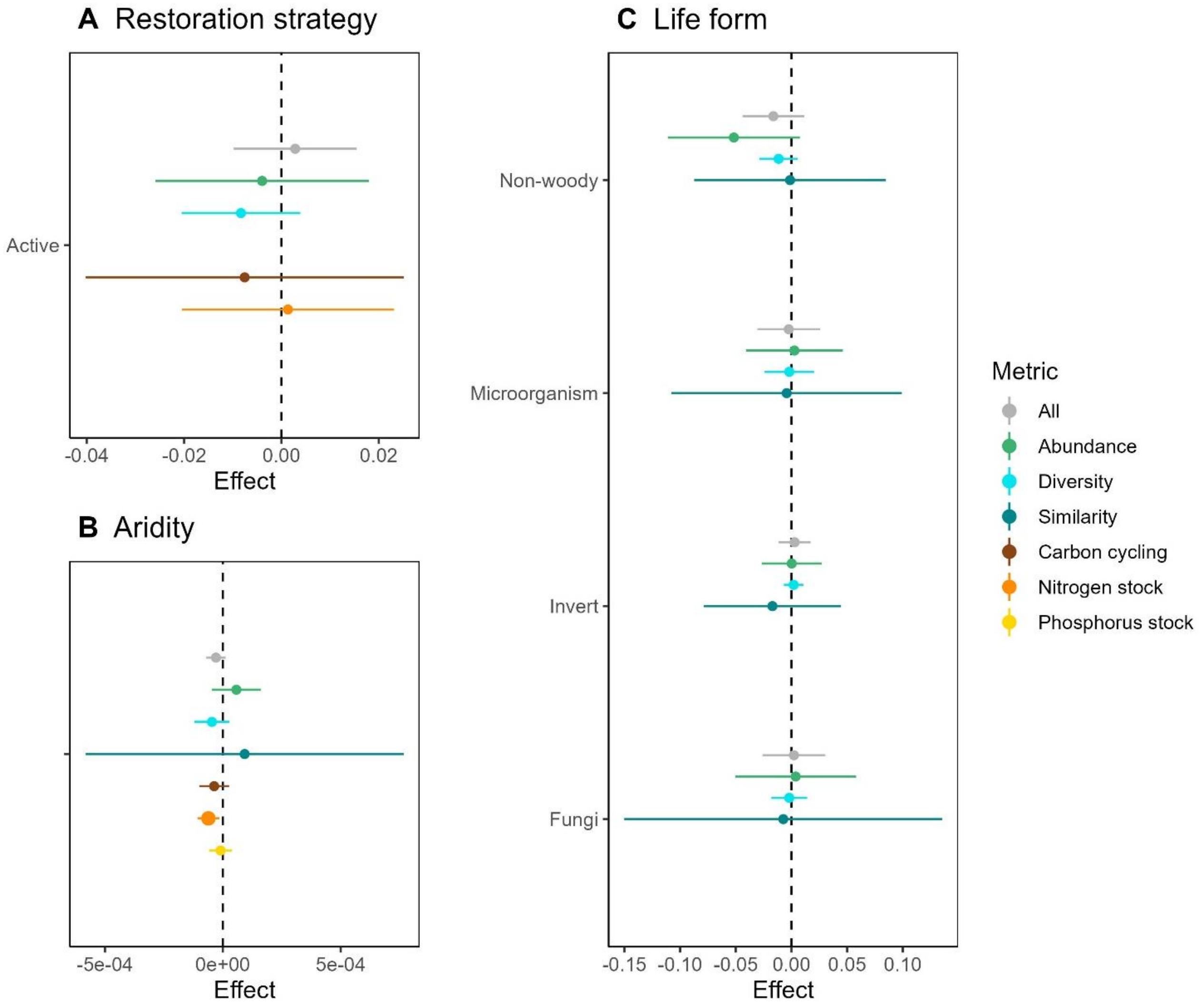


**Fig. S6.** Effect of the restoration strategy (A), aridity (B) and life form (C) on the recovery completeness of forest biodiversity (i.e., organism abundance, species diversity and Morisita-Horn species similarity) and functions (i.e., cycling of carbon, nitrogen stock and phosphorus stock), after 50 years since recovery started. Recovery completeness was computed with log response ratios. Error lines represent 95% confidence intervals of the estimated effect. Those error lines not overlapping the zero dashed line (shown with bigger dots) correspond to significant effects. The reference categories were “passive” (for restoration strategy) and “woody” (for life form). Having a positive effect means a higher recovery completeness with active restoration than with passive, and when the specific organism group is measured instead of woody plants.

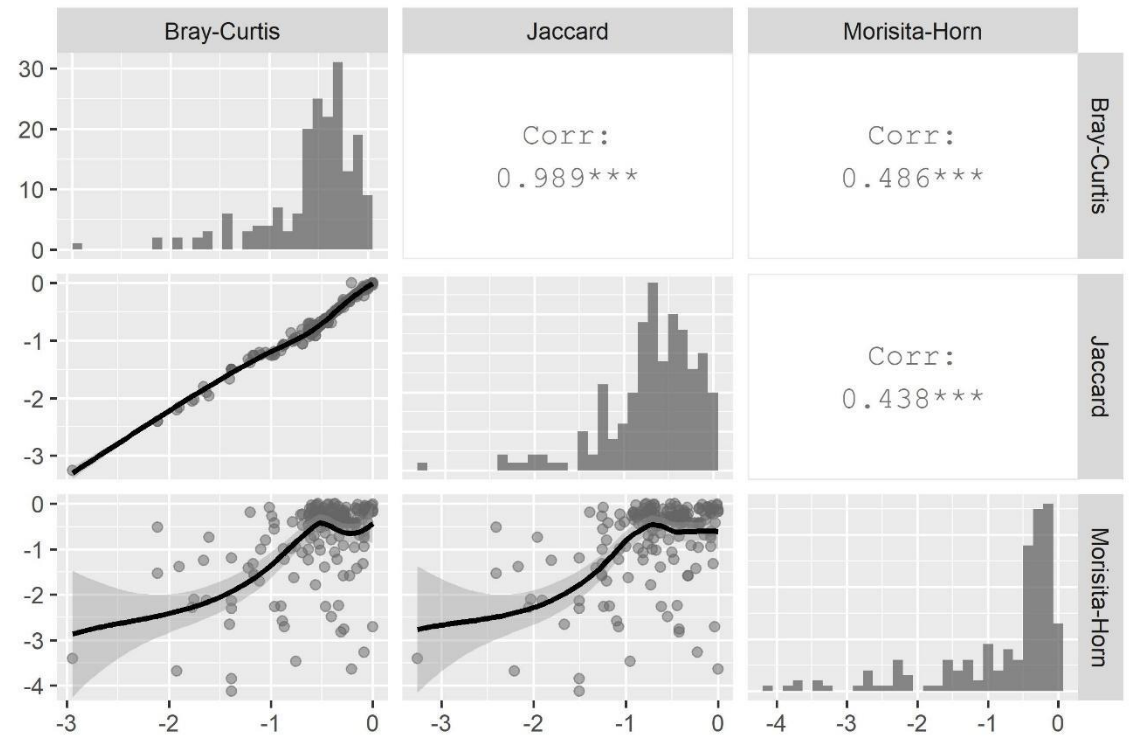


**Fig. S7.** Pairwise correlation analysis among response ratios for species similarity indices. Upper panels contain Pearson correlation index and the three asterisks indicate p-value is < 0.001; diagonal panels contains histograms of the response ratios for each similatiry index; lower panels are scatter plots with loess curves indicating the trend.

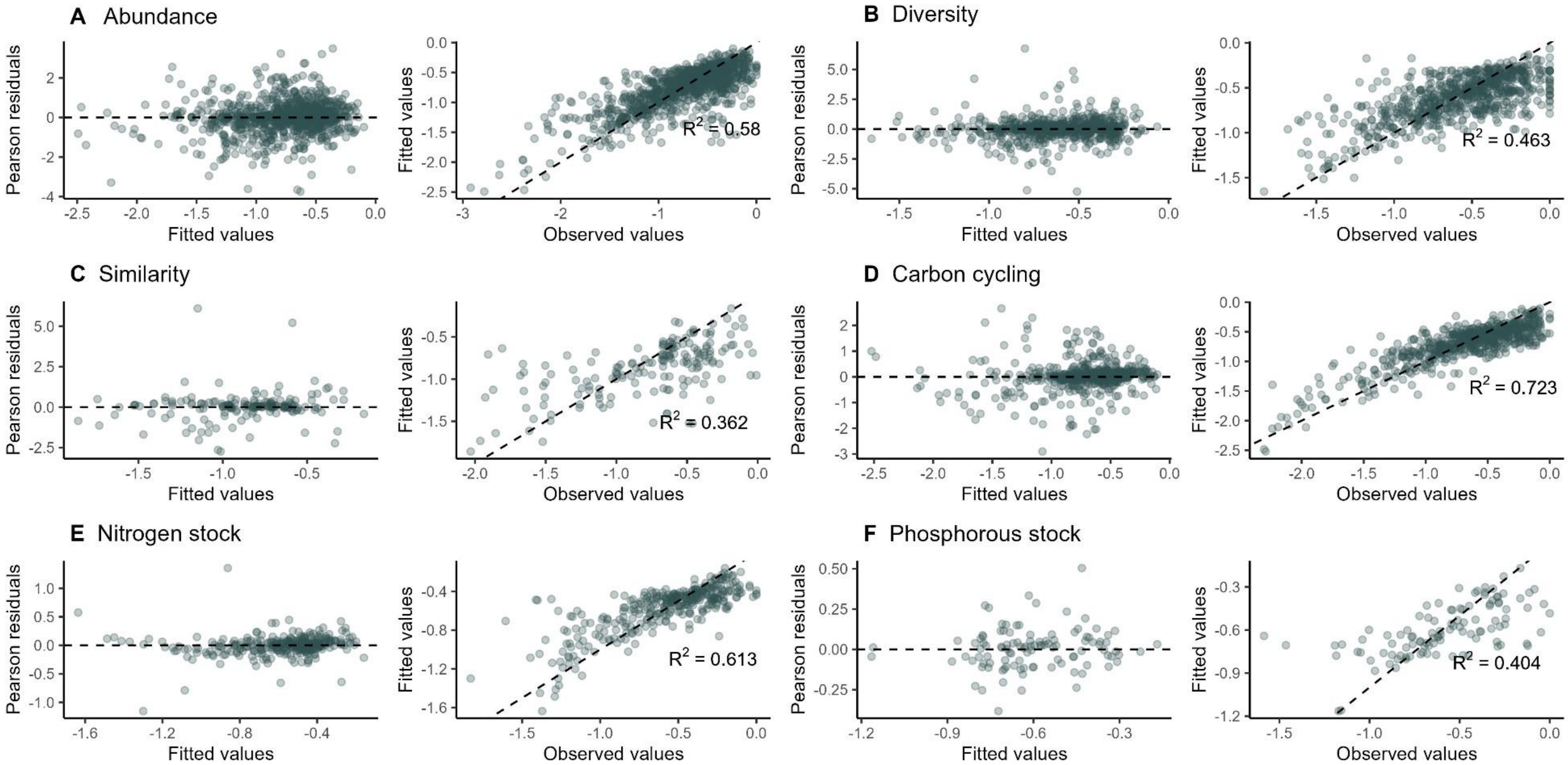


**Fig. S8.** Pearson residuals (left) and fitted vs. observed values (right) of the linear mixed models for the effect of recovery time on the response ratio of (A) organism abundance, (B) species diversity, (C) species similarity (Morisita-Horn index), (D) carbon cycling, (E) nitrogen stock and (F) phosphorus stock. Dashed lines represent data perfectly fitted to the model (Pearson residuals = 0) in left panels and 1:1 relationship in right panels.  The determination coefficient ($R^2$) of a linear regression between observed values and fitted values is also shown in left panels.

**Table S1.** Studies included in our meta-analysis database.

| Study | No. of chrono-sequences | Restoration strategy | Disturbance type | Recovery metric | | | | | |
|---|---|---|---|---|---|---|---|---|---|
| | | | | Abundance | Diversity | Similarity | Carbon | Nitrogen | Phosphorus |
| Addison et al. 2003 (*77*) | 1 | passive | logging | yes | yes | | | | |
| Addo-Fordjour et al. 2020 (*878* | 1 | passive | logging | yes | yes | yes | | | |
| Ashwood et al. 2019 (*79*) | 1 | active | cultivation | yes | | yes | yes | yes | |
| Aubin et al. 2009 (*80*) | 1 | passive | grazing | yes | yes | yes | yes | | |
| Batterman et al. 2013 (*81*) | 1 | passive | grazing | yes | | | yes | yes | |
| Bautista-Cruz et al. 2012 (*82*) | 1 | passive | cultivation | | | | yes | yes | |
| Becknell & Powers 2014 (*83*) | 1 | passive | agriculture | yes | yes | | yes | yes | yes |
| Bose et al. 2014 (*84*) | 1 | passive | cultivation | yes | yes | yes | yes | yes | yes |
| Bose et al. 2014 (*84*) | 2 | passive | cultivation | yes | yes | yes | yes | yes | yes |
| Bose et al. 2014 (*84*) | 3 | passive | cultivation | yes | yes | yes | yes | yes | yes |
| Bruelheide et al. 2011 (*85*) | 1 | passive | logging | | yes | yes | | | |
| Bu et al. 2014 (*86*) | 1 | passive | cultivation | yes | yes | | | | |
| Bu et al. 2014 (*87*) | 1 | passive | cultivation | | yes | | | | |
| Buzzard et al. 2016 (*88*) | 1 | passive | agriculture & logging | yes | yes | | yes | yes | yes |
| Chan et al. 2016 (*89*) | 1 | passive | cultivation | yes | yes | yes | yes | | |
| Ciarkowska 2017 (*90*) | 1 | passive | mining | | | | yes | | |
| Compton et al. 2007 (*91*) | 1 | passive | agriculture | | | | | yes | |
| Dalling & Denslow 1998 (*92*) | 1 | passive | agriculture | yes | yes | yes | | | |
| Davidson et al. 2007 (*93*) | 1 | passive | cultivation | yes | yes | yes | yes | yes | yes |
| Davidson et al. 2007 (*93*) | 2 | passive | cultivation | yes | yes | yes | yes | yes | yes |
| Deng et al. 2013 (*94*) | 1 | passive | cultivation | yes | | | yes | yes | |
| Deng et al. 2016 (*95*) | 1 | passive | cultivation | | | | yes | yes | |
| Deng et al. 2018 (*96*) | 1 | passive | cultivation | yes | | | yes | | |
| Denslow & Guzman 2000 (*97*) | 1 | passive | agriculture | yes | | | yes | | |
| Dent et al. 2013 (*98*) | 1 | passive | agriculture | | yes | yes | | | |
| DeWalt et al. 2003 (*99*) | 1 | passive | agriculture | yes | yes | yes | | | |
| Dungan et al. 2001 (*100*) | 1 | passive | cultivation | yes | yes | yes | | | |

| | | | | | | | | | |
|---|---|---|---|---|---|---|---|---|---|
| Foote & Grogan 2010 (*101*) | 1 | passive | cultivation | yes | yes | yes | yes | yes | yes |
| Foote & Grogan 2010 (*101*) | 2 | passive | cultivation | yes | yes | yes | yes | yes | yes |
| Foote & Grogan 2010 (*101*) | 3 | passive | cultivation | yes | yes | yes | yes | yes | yes |
| Fuss et al. 2019 (*102*) | 1 | passive | logging | | | | yes | yes | |
| Garcia et al. 2016 (*103*) | 1 | active | cultivation | yes | yes | | | | |
| Gießelmann et al. 2011 (*104*) | 1 | passive | grazing | | yes | | yes | | |
| Goebel & Hix 1996 (*105*) | 1 | passive | logging | yes | yes | yes | yes | | |
| Goosem et al. 2016 (*106*) | 1 | passive | grazing | | yes | | | | |
| Greenwood & Buttle 2014 (*107*) | 1 | active | agriculture | yes | | | yes | | |
| Hasegawa et al. 2006 (*108*) | 1 | passive | logging | yes | yes | | yes | | |
| Hasegawa et al. 2013 (*109*) | 1 | passive | logging | yes | yes | | | | |
| Herrero et al. 2016 (*110*) | 1 | passive | logging | | | | yes | | |
| Hingston & Grove 2009 (*111*) | 1 | passive | logging | | yes | | | | |
| Hopp et al. 2010 (*112*) | 1 | passive | agriculture & logging | yes | yes | yes | yes | yes | yes |
| Hopp et al. 2010 (*113*) | 2 | passive | agriculture & logging | yes | yes | yes | yes | yes | yes |
| Itioka et al. 2015 (*113*) | 1 | passive | cultivation | yes | yes | yes | | | |
| Janisch & Harmon 2002 (*114*) | 1 | passive | logging | | | | yes | | |
| Jones et al. 2019 (*115*) | 1 | passive | cultivation | yes | | | yes | | |
| Kahmen & Jules 2005 (*116*) | 1 | passive | logging | yes | | | | | |
| Kalinina et al. 2009 (*117*) | 1 | passive | cultivation | | | | yes | yes | |
| Kalinina et al. 2010 (*118*) | 1 | passive | cultivation | | | | yes | yes | |
| Kalinina et al. 2011 (*119*) | 1 | passive | grazing | | | | yes | yes | |
| Kalinina et al. 2015 (*120*) | 1 | passive | cultivation | yes | yes | yes | yes | yes | yes |
| Kalinina et al. 2015 (*120*) | 2 | passive | cultivation | yes | yes | yes | yes | yes | yes |
| Kalinina et al. 2015 (*120*) | 3 | passive | cultivation | yes | yes | yes | yes | yes | yes |
| Kalinina et al. 2018 (*121*) | 1 | passive | cultivation | | | | yes | | |
| Karelin et al. 2017 (*122*) | 1 | passive | cultivation | | | | yes | yes | |
| Kubota et al. 2005 (*123*) | 1 | passive | logging | yes | yes | | yes | yes | |
| Leiva et al. 2009 (*124*) | 1 | passive | grazing | yes | yes | yes | yes | yes | yes |
| Leiva et al. 2009 (*124*) | 2 | passive | grazing | yes | yes | yes | yes | yes | yes |

| | | | | | | | | | |
|---|---|---|---|---|---|---|---|---|---|
| Li et al. 2005 (*125*) | 1 | passive | cultivation | | | | yes | yes | |
| Liu et al. 2019 (*126*) | 1 | passive | logging | yes | | | yes | yes | yes |
| Long et al. 2012 (*127*) | 1 | passive | logging | | yes | | yes | yes | |
| López-Jiménez et al. 2019 (*128*) | 1 | passive | grazing | yes | yes | yes | | | |
| Lu et al. 2014 (*129*) | 1 | passive | cultivation | yes | yes | yes | | yes | yes |
| Lu et al. 2016 (*130*) | 1 | passive | cultivation | yes | yes | | yes | yes | yes |
| Lu et al. 2018 (*131*) | 1 | passive | cultivation | yes | | | yes | yes | yes |
| Lucas-Borja et al. 2019 (*132*) | 1 | passive | logging | yes | | | yes | yes | |
| Maeto & Sato 2004 (*133*) | 1 | passive | logging | yes | yes | yes | | | |
| Marín-Spiotta et al. 2007 (*134*) | 1 | passive | grazing | yes | yes | yes | | | |
| Marin-Spiotta et al. 2009 (*135*) | 1 | passive | grazing | | | | yes | yes | |
| Matlack 2009 (*136*) | 1 | passive | cultivation | yes | | | yes | | yes |
| McMahon et al. 2010 (*137*) | 1 | passive | agriculture & logging | yes | | | | | |
| McPherson & Timmer 2002 (*138*) | 1 | active | cultivation | yes | yes | yes | yes | yes | yes |
| McPherson & Timmer 2002 (*138*) | 2 | active | cultivation | yes | yes | yes | yes | yes | yes |
| Mi et al. 2016 (*139*) | 1 | passive | logging | yes | yes | | | yes | yes |
| Moola & Vasseur 2004 (*140*) | 1 | passive | logging | yes | yes | yes | | | |
| Moola & Vasseur 2009 (*141*) | 1 | passive | logging | yes | | | | | |
| Muñiz-Castro et al. 2006 (*142*) | 1 | passive | grazing | yes | yes | | | | |
| Muñiz-Castro et al. 2012 (*143*) | 1 | passive | grazing | yes | yes | yes | | | |
| Muñoz Gutiérrez et al. 2017 (*144*) | 1 | passive | cultivation | | yes | | | | |
| Negrete-Yankelevich et al. 2007 (*145*) | 1 | passive | logging | yes | yes | yes | yes | | |
| Osono & Trofymow 2012 (*146*) | 1 | active | logging | yes | yes | yes | yes | yes | yes |
| Osono & Trofymow 2012 (*146*) | 2 | active | logging | yes | yes | yes | yes | yes | yes |
| Ostertag et al. 2008 (*147*) | 1 | passive | grazing | | | | yes | yes | |
| Ottermanns et al. 2011 (*148*) | 1 | active | agriculture & logging | yes | | | yes | yes | |
| Panesar et al. 2001 (*149*) | 1 | passive | logging | yes | yes | | | | |
| Paz et al. 2016 (*150*) | 1 | passive | grazing | yes | yes | yes | yes | yes | yes |
| Paz et al. 2016 (*150*) | 2 | passive | grazing | yes | yes | yes | yes | yes | yes |
| Plue et al. 2010 (*151*) | 1 | passive | logging | yes | yes | yes | yes | yes | yes |

| | | | | | | | | | |
|---|---|---|---|---|---|---|---|---|---|
| Plue et al. 2010 (*151*) | 2 | passive | logging | yes | yes | yes | yes | yes | yes |
| Preston & Trofymow 2000 (*152*) | 1 | passive | logging | | | | yes | yes | yes |
| Preston et al. 2002 (*153*) | 1 | passive | logging | | | | yes | yes | |
| Ramos-Fabiel et al. 2019 (*154*) | 1 | passive | cultivation | yes | yes | yes | | | |
| Redi et al. 2005 (*155*) | 1 | passive | cultivation | yes | yes | yes | | | |
| Richardson et al. 2017 (*156*) | 1 | passive | logging | yes | yes | yes | yes | yes | yes |
| Richardson et al. 2017 (*156*) | 2 | passive | logging | yes | yes | yes | yes | yes | yes |
| Scheuermann et al. 2018 (*157*) | 1 | passive | logging | yes | yes | | | | |
| Schmidt et al. 2013 (*158*) | 1 | passive | cultivation | | yes | | | | |
| Serong & Lill 2008 (*159*) | 1 | passive | logging | yes | yes | | | | |
| Shao et al. 2019 (*160*) | 1 | passive | logging | | | | yes | yes | |
| Shoo et al. 2016 (*161*) | 1 | passive | agriculture | yes | yes | | | | |
| Simmons et al. 2015 (*162*) | 1 | passive | cultivation | yes | | | | | |
| Smith et al. 2015 (*163*) | 1 | passive | grazing | yes | | | | | |
| Sturtevant et al. 1997 (*164*) | 1 | passive | logging | yes | | | yes | | |
| Suganuma & Durigan 2015 (*165*) | 1 | active | agriculture | yes | yes | | | | |
| Sutherland et al. 2016 (*166*) | 1 | passive | logging | yes | | | yes | | |
| Tabarelli & Peres 2002 (*167*) | 1 | passive | cultivation | | yes | | | | |
| Tang et al. 2009 (*168*) | 1 | passive | logging | yes | | | yes | | |
| Tausan et al. 2017 (*169*) | 1 | passive | logging | yes | yes | yes | yes | yes | yes |
| Tausan et al. 2017 (*169*) | 2 | passive | logging | yes | yes | yes | yes | yes | yes |
| Thuille et al. 2000 (*170*) | 1 | passive | grazing | yes | | | yes | | |
| Van Gemerden et al. 2003 (*171*) | 1 | passive | cultivation | yes | yes | | | | |
| Wang et al. 2016 (*172*) | 1 | passive | cultivation | yes | | | yes | | |
| Williams & Moriarity 2000 (*173*) | 1 | passive | logging | yes | yes | yes | yes | | |
| Winbourne et al. 2018 (*174*) | 1 | passive | cultivation | yes | | | yes | yes | yes |
| Woods & DeWalt 2013 (*175*) | 1 | passive | agriculture | yes | yes | | | | |
| Yamashita et al. 2012 (*176*) | 1 | passive | logging | yes | yes | | yes | | |
| Yan et al. 2007 (*177*) | 1 | passive | logging | | yes | | yes | | |
| Zhang et al. 2010 (*178*) | 1 | passive | cultivation | yes | | | yes | yes | yes |

| | | | | | | | | | |
|---|---|---|---|---|---|---|---|---|---|
| Zhang et al. 2011 (*179*) | 1 | active | agriculture & logging | yes | yes | | yes | yes | |
| Zhang & Shangguan 2016 (*180*) | 1 | passive | cultivation | yes | | | yes | | |
| Zhang et al. 2018 (*181*) | 1 | passive | cultivation | | | | yes | yes | yes |
| Zhao et al. 2015 (*182*) | 1 | passive | cultivation | yes | | | yes | yes | |
| Zheng et al. 2019 (*183*) | 1 | passive | logging | yes | | | yes | yes | |
| Zhu et al. 2009 (*184*) | 1 | passive | logging | | yes | yes | | | |
| Zhu et al. 2010a (*185*) | 1 | passive | logging | yes | yes | yes | yes | yes | yes |
| Zhu et al. 2010b (*186*) | 1 | passive | cultivation | yes | yes | yes | yes | yes | yes |

**Table S2.** Distribution of primary studies, chronosequences, trajectories, data-points, time since recovery started (i.e. recovery time), reference sites and study areas by disturbance type, restoration strategy, recovery metric and life form studied. The distribution by life form studied only includes organism abundance, species diversity and species similarity trajectories. For the distribution by metric, the sum of the number of studies, chronosequences, reference sites and study areas is higher than for disturbance type and restoration strategy, because the metric categories are not exclusive.

| **Disturbance** | **No. studies** | **No. chrono-sequences** | **No. trajectories** | **No. data-points** | **Average (min. - max.) no. data-points per trajectory** | **Average (min. - max.) recovery time of each trajectory** | **No. reference sites** | **Area of the study site (km$^2$)** | **Studies reporting restored area (%)** |
|---|---|---|---|---|---|---|---|---|---|
| Agriculture | 10 | 13 | 59 | 880 | 15 (8 - 39) | 90 (53 - 115) | 15 | 3,021 | 90 |
| Agriculture & logging | 5 | 6 | 26 | 230 | 9 (2 - 54) | 96 (50 - 200) | 8 | 1,826 | 100 |
| Cultivation | 40 | 42 | 221 | 1413 | 7 (2 - 34) | 75 (50 - 170) | 116 | 126,900 | 90 |
| Logging | 39 | 43 | 243 | 2658 | 11 (2 - 34) | 96 (50 - 295) | 89 | 49,901 | 97 |
| Mining | 1 | 1 | 2 | 20 | 10 | 100 | 2 | 0.0012 | 100 |
| Grazing | 15 | 17 | 90 | 794 | 9 (4 - 33) | 69 (50 - 90) | 22 | 1,852 | 80 |
| **Restoration** | | | | | | | | | |
| Passive | 102 | 112 | 567 | 5361 | 10 (2 - 54) | 85 (50 - 295) | 231 | 181,731 | 89 |
| Active | 8 | 10 | 74 | 634 | 9 (3 - 29) | 83 (50 - 143) | 22 | 1,770 | 100 |
| **Metric** | | | | | | | | | |
| Abundance | 74 | 81 | 209 | 2001 | 10 (2 - 54) | 85 (50 - 295) | 160 | 128,224 | 95 |
| Diversity | 56 | 60 | 160 | 1411 | 9 (2 - 39) | 80 (50 - 150) | 86 | 21,901 | 98 |
| Similarity | 28 | 31 | 42 | 310 | 8 (2 - 34) | 87 (50 - 150) | 47 | 13,226 | 96 |
| Carbon | 60 | 66 | 128 | 1338 | 10 (2 - 39) | 89 (50 - 200) | 168 | 170,846 | 93 |
| Nitrogen | 37 | 38 | 77 | 711 | 9 (2 - 39) | 88 (50 - 200) | 66 | 100,390 | 89 |
| Phosphorus | 15 | 15 | 25 | 224 | 9 (2 - 39) | 71 (50 - 110) | 58 | 26,770 | 100 |
| **Life form** | | | | | | | | | |
| Bird | 1 | 1 | 2 | 18 | 9 | 111 | 1 | 69 | 100 |
| Fungi | 5 | 6 | 15 | 84 | 6 (4 - 8) | 94 (50 - 129) | 8 | 50 | 80 |
| Invertebrate | 15 | 17 | 72 | 695 | 10 (2 - 34) | 83 (54 - 113) | 37 | 4,710 | 93 |
| Microorganism | 4 | 4 | 17 | 95 | 5 (3 - 6) | 86 (50 - 100) | 5 | 260 | 75 |
| Non-woody | 7 | 7 | 18 | 133 | 7 (2 - 19) | 131 (55 - 295) | 8 | 5,578 | 100 |
| Woody | 61 | 67 | 254 | 2938 | 10 (2 - 54) | 81 (50 - 200) | 140 | 80,066 | 92 |
| Woody & non-woody | 14 | 14 | 33 | 170 | 5 (3 - 28) | 69 (53 - 130) | 17 | 54,200 | 100 |

**Table S3.** Results of the models fitted for the response ratio (RR) of each recovery metric. The response variable of the models equals the sqrt-transformed absolute RRs multiplied by -1. SE: standard error. Bold p-values were considered significant.

| Organism abundance | | | | |
|---|---|---|---|---|
| **Predictor variable** | **Estimate** | **SE** | ***t*-statistic** | ***p*-value** |
| Intercept | -0.76 | 0.03 | -24.51 | **<0.001** |
| ln(recovery time + 1) | 0.27 | 0.03 | 10.43 | **<0.001** |
| group trajectory identity - sd (intercept) | 0.34 | | | |
| group trajectory identity - sd (ln(recovery time + 1)) | 0.25 | | | |
| group trajectory identity - Correlation: intercept & ln(recovery time + 1) | -0.56 | | | |
| sd (residual) | 0.88 | | | |
| Marginal $R^2$ (fixed factors) | 0.07 | | | |
| Conditional $R^2$ (fixed and random factors) | 0.25 | | | |

| Species diversity | | | | |
|---|---|---|---|---|
| **Predictor variable** | **Estimate** | **SE** | ***t*-statistic** | ***p*-value** |
| Intercept | -0.62 | 0.03 | -23.13 | **<0.001** |
| √recovery time | 0.17 | 0.02 | 10.45 | **<0.001** |
| group trajectory identity - sd (intercept) | 0.24 | | | |
| group trajectory identity - sd (√recovery time) | 0.12 | | | |
| group trajectory identity - Correlation: intercept & √recovery time | -0.75 | | | |
| sd (residual) | 0.95 | | | |
| Marginal $R^2$ (fixed factors) | 0.03 | | | |
| Conditional $R^2$ (fixed and random factors) | 0.10 | | | |

| Morisita-Horn similarity | | | | |
|---|---|---|---|---|
| **Predictor variable** | **Estimate** | **SE** | ***t*-statistic** | ***p*-value** |
| Intercept | -0.90 | 0.10 | -9.27 | **<0.001** |
| ln(recovery time + 1) | 0.23 | 0.08 | 2.83 | **0.010** |
| group trajectory identity - sd (intercept) | 0.42 | | | |
| group trajectory identity - sd (ln(recovery time + 1)) | 0.31 | | | |
| group trajectory identity - Correlation: intercept & ln(recovery time + 1) | -0.58 | | | |
| sd (residual) | 1.02 | | | |
| Marginal $R^2$ (fixed factors) | 0.04 | | | |

| | | | | |
|---|---|---|---|---|
| Conditional $R^2$ (fixed and random factors) | 0.24 | | | |
| **Carbon cycling** | | | | |
| **Predictor variable** | **Estimate** | **SE** | ***t*-statistic** | ***p*-value** |
| Intercept | -0.64 | 0.04 | -15.29 | **<0.001** |
| √recovery time | 0.17 | 0.03 | 5.48 | **<0.001** |
| group trajectory identity - sd (intercept) | 0.34 | | | |
| group trajectory identity - sd (√recovery time) | 0.24 | | | |
| group trajectory identity - Correlation: intercept & √recovery time | -0.68 | | | |
| sd (residual) | 0.59 | | | |
| Marginal $R^2$ (fixed factors) | 0.06 | | | |
| Conditional $R^2$ (fixed and random factors) | 0.37 | | | |
| **Nitrogen stock** | | | | |
| **Predictor variable** | **Estimate** | **SE** | ***t*-statistic** | ***p*-value** |
| Intercept | -0.54 | 0.03 | -16.93 | **<0.001** |
| ln(recovery time + 1) | 0.11 | 0.02 | 5.07 | **<0.001** |
| group trajectory identity - sd (intercept) | 0.20 | | | |
| group trajectory identity - sd (ln(recovery time + 1)) | 0.11 | | | |
| group trajectory identity - Correlation: intercept & ln(recovery time + 1) | -0.51 | | | |
| sd (residual) | 0.19 | | | |
| Marginal $R^2$ (fixed factors) | 0.12 | | | |
| Conditional $R^2$ (fixed and random factors) | 0.65 | | | |
| **Phosphorus stock** | | | | |
| **Predictor variable** | **Estimate** | **SE** | ***t*-statistic** | ***p*-value** |
| Intercept | -0.61 | 0.05 | -11.51 | **<0.001** |
| ln(recovery time + 1) | 0.06 | 0.02 | 3.38 | **0.005** |
| group trajectory identity - sd (intercept) | 0.23 | | | |
| group trajectory identity - sd (ln(recovery time + 1)) | 0.28 | | | |
| group trajectory identity - Correlation: intercept & ln(recovery time + 1) | 0.14 | | | |
| sd (residual) | 0.05 | | | |
| Marginal $R^2$ (fixed factors) | 0.05 | | | |
| Conditional $R^2$ (fixed and random factors) | 0.74 | | | |

**Table S4.** Results of the liner models fitted for the effect of the recovery metric on the predicted response ratio after 50 and 100 years of recovery. The reference level for the recovery metric is organism abundance. Bold p-values were considered significant.

| | **50 years** | | | |
|---|---|---|---|---|
| **Predictor variable** | **Estimate** | **SE** | ***t*-statistic** | ***p*-value** |
| (LMM intercept) | -0.03 | 0.007 | -4.3 | **<0.001** |
| Diversity | 0.02 | 0.005 | 3.24 | **0.002** |
| Similarity | -0.02 | 0.008 | -2.80 | **0.036** |
| Carbon | -0.02 | 0.006 | -3.22 | **0.001** |
| Nitrogen | -0.01 | 0.007 | -1.65 | 0.094 |
| Phosphorus | -0.05 | 0.010 | -4.41 | **<0.001** |
| Intercept | 0.81 | 0.009 | 94.76 | **<0.001** |
| $R^2$ | 0.95 | | | |

| | **100 years** | | | |
|---|---|---|---|---|
| **Predictor variable** | **Estimate** | **SE** | ***t*-statistic** | ***p*-value** |
| (LMM intercept) | -0.147 | 0.03 | -5.00 | **<0.001** |
| Diversity | 0.096 | 0.02 | 3.93 | **<0.001** |
| Similarity | -0.093 | 0.03 | -2.97 | **0.003** |
| Carbon | -0.005 | 0.02 | -0.22 | 0.824 |
| Nitrogen | 0.007 | 0.03 | 0.25 | 0.803 |
| Phosphorus | 0.032 | 0.07 | 0.44 | 0.658 |
| Intercept | 0.466 | 0.03 | 14.99 | **<0.001** |
| $R^2$ | 0.65 | | | |

**Table S5.** Comparisons of the three models fitted to test a linear, logarithmic or square root effect of the recovery time on the response ratio of each metric. Df: degrees of freedom. we consider three functions to include the recovery time variableA ICc: Akaike Information Criterion corrected for small samples. ΔAICc: AICc increase compared to the best model. The best model is shown in bold.

| Metric type | linear | | | ln(recovery time + 1) | | | √recovery time | | |
|---|---|---|---|---|---|---|---|---|---|
| | **AICc** | **ΔAICc** | **Df** | **AICc** | **ΔAICc** | **Df** | **AICc** | **ΔAICc** | **Df** |
| Abundance | 2212.49 | 217.42 | 6 | **1995.07** | **0** | **6** | 2059.41 | 64.34 | 6 |
| Diversity | 1688.06 | 36.27 | 6 | 1651.79 | 10.59 | 6 | **1641.20** | **0** | **6** |
| Similarity | 526.98 | 23.93 | 6 | **503.05** | **0** | **6** | 511.83 | 8.78 | 6 |
| Carbon | 1435.35 | 9.23 | 6 | 1426.12 | 48.33 | 6 | **1377.79** | **0** | **6** |
| Nitrogen | 468.27 | 69.46 | 6 | **398.81** | **0** | **6** | 406.39 | 7.58 | 6 |
| Phosphorus | **53.88** | **0** | **6** | 61.42 | 7.54 | 6 | 55.64 | 1.76 | 6 |

**Table S6.** Pairwise comparisons among the categories of the variable "recovery metric" from the models fitted for the effect of the recovery metric on the predicted response ratio after 50 and 100 years of recovery. Bold p-values were considered significant.

| | | | | | | |
|---|---|---|---|---|---|---|
| | | | **50 years** | | | |
| **Comparisons** | **Estimate** | **SE** | ***t*-ratio** | ***p*-value** | **value ± SE** | **CI (0.95)** |
| Abundance | | | | | -0.57 ± 0.003 | [-0.58, -0.57] |
| Abundance – Diversity | -0.02 | 0.005 | -3.24 | **0.016** | | |
| Abundance – Similarity | 0.02 | 0.009 | 2.09 | 0.292 | | |
| Abundance – Carbon | 0.02 | 0.006 | 3.22 | **0.017** | | |
| Abundance – Nitrogen | 0.01 | 0.006 | 1.68 | 0.558 | | |
| Abundance – Phosphorus | 0.05 | 0.010 | 4.39 | **<0.001** | | |
| Diversity | | | | | -0.56 ± 0.004 | [-0.56, -0.55] |
| Diversity – Similarity | 0.04 | 0.009 | 3.82 | **0.002** | | |
| Diversity – Carbon | 0.03 | 0.006 | 6.00 | **<0.001** | | |
| Diversity – Nitrogen | 0.03 | 0.007 | 4.19 | **<0.001** | | |
| Diversity – Phosphorus | 0.06 | 0.010 | 5.97 | **<0.001** | | |
| Similarity | | | | | -0.59 ± 0.008 | [-0.61, -0.58] |
| Similarity – Carbon | -0.0004 | 0.009 | -0.05 | 1.000 | | |
| Similarity – Nitrogen | -0.007 | 0.010 | -0.71 | 0.981 | | |
| Similarity – Phosphorus | 0.027 | 0.013 | 2.10 | 0.289 | | |
| Carbon | | | | | -0.59 ± 0.004 | [-0.60, -0.58] |
| Carbon – Nitrogen | -0.007 | 0.007 | -0.97 | 0.927 | | |
| Carbon – Phosphorus | 0.027 | 0.010 | 2.58 | 0.103 | | |
| Nitrogen | | | | | -0.58 ± 0.005 | [-0.59, -0.57] |
| Nitrogen – Phosphorus | 0.03 | 0.011 | 3.07 | **0.027** | | |
| Phosphorus | | | | | -0.62 ± 0.010 | [-0.64, -0.60] |
| | | | **100 years** | | | |
| **Comparisons** | **Estimate** | **SE** | ***t*-ratio** | ***p*-value** | **value ± SE** | **CI (0.95)** |
| Abundance | | | | | -0.47 ± 0.02 | [-0.50, -0.44] |
| Abundance – Diversity | -0.096 | 0.02 | -3.91 | **0.002** | | |
| Abundance – Similarity | 0.084 | 0.03 | 2.69 | 0.082 | | |
| Abundance – Carbon | 0.006 | 0.02 | 0.23 | 1.000 | | |
| Abundance – Nitrogen | -0.007 | 0.03 | -0.23 | 1.000 | | |
| Abundance – Phosphorus | -0.031 | 0.07 | -0.44 | 0.998 | | |
| Diversity | | | | | -0.37 ± 0.02 | [-0.41, -0.34] |
| Diversity – Similarity | 0.18 | 0.03 | 5.30 | **<0.001** | | |
| Diversity – Carbon | 0.10 | 0.03 | 4.03 | **0.001** | | |
| Diversity – Nitrogen | 0.09 | 0.03 | 3.03 | **0.033** | | |
| Diversity – Phosphorus | 0.06 | 0.07 | 0.89 | 0.948 | | |
| Similarity | | | | | -0.55 ± 0.03 | [-0.61, -0.50] |
| Similarity – Carbon | -0.08 | 0.03 | -2.63 | 0.175 | | |
| Similarity – Nitrogen | -0.09 | 0.04 | -2.41 | 0.158 | | |
| Similarity – Phosphorus | -0.12 | 0.08 | -1.53 | 0.645 | | |
| Carbon | | | | | -0.47 ± 0.02 | [-0.51, -0.44] |
| Carbon – Nitrogen | -0.01 | 0.03 | -0.42 | 0.998 | | |

| | | | | | | |
|---|---|---|---|---|---|---|
| Carbon – Phosphorus | -0.04 | 0.07 | -0.51 | 0.995 | | |
| Nitrogen | | | | | -0.46 ± 0.02 | [-0.51, -0.41] |
| Nitrogen – Phosphorus | -0.02 | 0.07 | -0.33 | 1.000 | | |
| Phosphorus | | | | | -0.44 ± 0.07 | [-0.58, -0.30] |

**Table S7.** Percentage of recovery of each recovery metric at different times since recovery started: 73, 146 and 219 years of recovery (i.e., one, two and three times the global life expectancy). CI: Confidence Interval.

| | **73 years** | | **146 years** | | **219 years** | |
|---|---|---|---|---|---|---|
| **Recovery metric** | **Recovery completeness (%)** | **CI (0.95)** | **Recovery completeness (%)** | **CI (0.95)** | **Recovery completeness (%)** | **CI (0.95)** |
| Abundance | 74.78 | [70.36 – 79.02] | 88.43 | [83.76 – 92.45] | 94.46 | [90.01 – 97.69] |
| Diversity | 84.45 | [81.25 – 87.43] | 97.09 | [94.33 – 98.96] | | |
| Similarity | 61.34 | [48.03 – 74.56] | 73.18 | [56.33 – 87.85] | | |
| Carbon | 79.51 | [74.01 – 84.59] | 92.74 | [86.28 – 97.32] | | |
| Nitrogen | 82.55 | [78.16 – 86.60] | 87.24 | [82.48 – 91.40] | | |
| Phosphorus | 76.39 | [66.15 – 85.56] | | | | |

**Table S8.** Estimated time to reach 90% of reference values for each recovery metric based on a prediction for each trajectory.

| Metric type | Median time to recovery (years) | 10% quantile | 90% quantile |
|---|---|---|---|
| Abundance | 160.09 | 54.64 | 466.34 |
| Diversity | 96.59 | 64.06 | 121.88 |
| Similarity | 494.12 | 91.82 | 2039.27 |
| Carbon | 125.91 | 68.73 | 169.08 |
| Nitrogen | 218.48 | 37.58 | 745.12 |
| Phosphorus | 134.35 | 42.00 | 279.76 |

**Table S9.** Results of the linear models fitted for the effect of aridity, disturbance category, restoration strategy and life form on the response ratio predicted for all recovery metrics together, after 50 and 100 years of recovery. The reference level for the disturbance type is "logging", for the restoration strategy is "passive" and for the life form is "woody". Bold *p*-values were considered significant.

| | | | | | | | | |
|---|---|---|---|---|---|---|---|---|
| | ***Aridity*** | | | | | | | |
| | **50 years (n = 596)** | | | | **100 years (n = 202)** | | | |
| **Predictor variable** | **Estimate** | **SE** | ***t*-statistic** | ***p*-value** | **Estimate** | **SE** | ***t*-statistic** | ***p*-value** |
| (LMM intercept) | -0.03 | 0.007 | -4.60 | **<0.001** | -0.06 | 0.03 | -2.23 | **0.027** |
| Aridity | -0.00003 | 0.00002 | -1.40 | 0.165 | -0.0003 | 0.0001 | -2.57 | **0.011** |
| Intercept | 0.81 | 0.008 | 98.07 | **<0.001** | 0.51 | 0.03 | 16.97 | **<0.001** |
| $R^2$ | 0.95 | | | | 0.61 | | | |
| | ***Disturbance type*** | | | | | | | |
| | **50 years (n = 572)** | | | | **100 years (n = 179)** | | | |
| **Predictor variable** | **Estimate** | **SE** | ***t*-statistic** | ***p*-value** | **Estimate** | **SE** | ***t*-statistic** | ***p*-value** |
| (LMM intercept) | -0.04 | 0.006 | -6.63 | **<0.001** | -0.106 | 0.025 | -4.19 | **<0.001** |
| Agriculture | 0.007 | 0.007 | 0.95 | 0.344 | 0.006 | 0.024 | 0.27 | 0.788 |
| Cultivation | 0.009 | 0.005 | 1.92 | **0.055** | 0.034 | 0.022 | 1.53 | 0.129 |
| Grazing | 0.031 | 0.006 | 4.81 | **<0.001** | - | - | - | - |
| Intercept | 0.807 | 0.008 | 97.54 | **<0.001** | 0.519 | 0.031 | 16.55 | **<0.001** |
| $R^2$ | 0.94 | | | | 0.62 | | | |
| | ***Restoration strategy*** | | | | | | | |
| | **50 years (n = 596)** | | | | **100 years (n = 202)** | | | |
| **Predictor variable** | **Estimate** | **SE** | ***t*-statistic** | ***p*-value** | **Estimate** | **SE** | ***t*-statistic** | ***p*-value** |
| (LMM intercept) | -0.034 | 0.006 | -5.69 | **<0.001** | -0.10 | 0.02 | -4.15 | **<0.001** |
| Active | 0.003 | 0.006 | 0.44 | 0.661 | 0.02 | 0.03 | 0.61 | 0.545 |
| Intercept | 0.809 | 0.008 | 97.69 | **<0.001** | 0.52 | 0.03 | 17.28 | **<0.001** |
| $R^2$ | 0.94 | | | | 0.60 | | | |
| | ***Life form*** | | | | | | | |
| | **50 years (n = 342)** | | | | **100 years (n = 101)** | | | |
| **Predictor variable** | **Estimate** | **SE** | **t-statistic** | ***p*-value** | **Estimate** | **SE** | ***t*-statistic** | ***p*-value** |

| | | | | | | | | |
|---|---|---|---|---|---|---|---|---|
| (LMM intercept) | -0.014 | 0.009 | -1.61 | 0.107 | -0.043 | 0.030 | -1.42 | 0.158 |
| Fungi | 0.002 | 0.014 | 0.16 | 0.875 | 0.047 | 0.043 | 1.10 | 0.273 |
| Invertebrate | 0.003 | 0.007 | 0.40 | 0.693 | 0.035 | 0.035 | 1.00 | 0.319 |
| Microorganism | -0.002 | 0.014 | -0.17 | 0.866 | 0.029 | 0.041 | 0.71 | 0.482 |
| Non-woody | -0.016 | 0.014 | -1.15 | 0.250 | 0.010 | 0.041 | 0.25 | 0.805 |
| Intercept | 0.822 | 0.012 | 70.16 | **<0.001** | 0.593 | 0.038 | 15.80 | **<0.001** |
| $R^2$ | 0.94 | | | | 0.71 | | | |

**Table S10.** Pairwise comparisons among the categories of the variable "disturbance category" for their effect on the response ratio predicted for all recovery metrics together, after 50 years of recovery. Bold *p*-values were considered significant.

| 50 years | | | | | | |
|---|---|---|---|---|---|---|
| **Comparisons** | **Estimate** | **SE** | ***t*-ratio** | ***p*-value** | **value ± SE** | **CI (0.95)** |
| Logging | | | | | -0.583 ± 0.003 | [-0.59, -0.58] |
| Logging – Agriculture | -0.007 | 0.007 | -0.95 | 0.780 | | |
| Logging – Cultivation | -0.009 | 0.005 | -1.92 | 0.221 | | |
| Logging – Grazing | -0.031 | 0.006 | -4.81 | <**0.001** | | |
| Agriculture | | | | | -0.576 ± 0.007 | [-0.59, -0.56] |
| Agriculture – Cultivation | -0.002 | 0.007 | -0.26 | 0.994 | | |
| Agriculture – Grazing | -0.024 | 0.009 | -2.79 | **0.028** | | |
| Cultivation | | | | | -0.574 ± 0.003 | [-0.58, -0.57] |
| Cultivation – Grazing | -0.022 | 0.007 | -3.39 | **0.004** | | |
| Grazing | | | | | -0.552 ± 0.006 | [-0.56, -0.54] |

**Table S11.** Results of the linear models fitted for the effect of aridity, disturbance category, restoration strategy and life form on the response ratio predicted for organism abundance after 50 and 100 years of recovery. The reference level for the disturbance type is "logging", for the restoration strategy is "passive" and for the life form is "woody". Bold *p*-values were considered significant.

| ***Aridity*** | | | | | | | | |
|---|---|---|---|---|---|---|---|---|
| | **50 years (n = 195)** | | | | **100 years (n = 59)** | | | |
| **Predictor variable** | **Estimate** | **SE** | ***t*-statistic** | ***p*-value** | **Estimate** | **SE** | ***t*-statistic** | ***p*-value** |
| (LMM intercept) | -0.02 | 0.01 | -1.40 | 0.162 | -0.01 | 0.05 | -0.31 | 0.76 |
| Aridity | -0.00006 | 0.00005 | 1.08 | 0.279 | -0.0008 | 0.0002 | -3.35 | **0.001** |
| Intercept | 0.83 | 0.02 | 49.21 | **<0.001** | 0.50 | 0.05 | 10.53 | **<0.001** |
| $R^2$ | 0.93 | | | | 0.72 | | | |
| ***Disturbance type*** | | | | | | | | |
| | **50 years(n = 187)** | | | | **100 years (n = 51)** | | | |
| **Predictor variable** | **Estimate** | **SE** | ***t*-statistic** | ***p*-value** | **Estimate** | **SE** | ***t*-statistic** | ***p*-value** |
| (LMM intercept) | -0.028 | 0.016 | -1.80 | 0.073 | -0.113 | 0.05 | -2.15 | **0.037** |
| Agriculture | 0.003 | 0.018 | 0.24 | 0.811 | 0.021 | 0.05 | 0.46 | 0.651 |
| Cultivation | 0.007 | 0.011 | 0.66 | 0.509 | 0.080 | 0.06 | 1.40 | 0.166 |
| Grazing | 0.033 | 0.013 | 2.60 | **0.010** | - | - | - | - |
| Intercept | 0.823 | 0.017 | 48.37 | **<0.001** | 0.536 | 0.05 | 9.71 | **<0.001** |
| $R^2$ | 0.93 | | | | 0.66 | | | |
| ***Restoration strategy*** | | | | | | | | |
| | **50 years (n = 195)** | | | | **100 years (n = 59)** | | | |
| **Predictor variable** | **Estimate** | **SE** | ***t*-statistic** | ***p*-value** | **Estimate** | **SE** | ***t*-statistic** | ***p*-value** |
| (LMM intercept) | -0.015 | 0.01 | -1.13 | 0.259 | -0.095 | 0.05 | -2.11 | 0.039 |
| Active | -0.004 | 0.01 | -0.36 | 0.723 | 0.049 | 0.05 | 0.94 | 0.350 |
| Intercept | 0.827 | 0.02 | 50.89 | **<0.001** | 0.541 | 0.05 | 10.74 | **<0.001** |
| $R^2$ | 0.93 | | | | 0.67 | | | |
| ***Life form*** | | | | | | | | |
| | **50 years (n = 182)** | | | | **100 years (n = 55)** | | | |

| **Predictor variable** | **Estimate** | **SE** | **t-statistic** | ***p*-value** | **Estimate** | **SE** | ***t*-statistic** | ***p*-value** |
|---|---|---|---|---|---|---|---|---|
| (LMM intercept) | -0.030 | 0.016 | -1.82 | 0.070 | -0.17 | 0.06 | -2.77 | **0.008** |
| Fungi | 0.004 | 0.028 | 0.14 | 0.891 | 0.03 | 0.11 | 0.29 | 0.774 |
| Invertebrate | 0.0003 | 0.014 | 0.02 | 0.985 | 0.05 | 0.06 | 0.86 | 0.392 |
| Microorganism | 0.003 | 0.022 | 0.12 | 0.904 | 0.04 | 0.08 | 0.59 | 0.559 |
| Non-woody | -0.052 | 0.030 | -1.71 | 0.089 | -0.11 | 0.09 | -1.25 | 0.219 |
| Intercept | 0.806 | 0.021 | 39.00 | **<0.001** | 0.45 | 0.07 | 6.00 | **<0.001** |
| $R^2$ | 0.93 | | | | 0.64 | | | |

**Table S12.** Results of the linear models fitted for the effect of aridity, disturbance category, restoration strategy and life form on the response ratio predicted for species diversity after 50 and 100 years of recovery. The reference level for the disturbance type is "logging", for the restoration strategy is "passive" and for the life form is "woody". Bold *p*-values were considered significant.

***Aridity***

| | **50 years (n = 144)** | | | | **100 years (n = 49)** | | | |
|---|---|---|---|---|---|---|---|---|
| **Predictor variable** | **Estimate** | **SE** | ***t*-statistic** | ***p*-value** | **Estimate** | **SE** | ***t*-statistic** | ***p*-value** |
| (LMM intercept) | -0.04 | 0.01 | -4.21 | **<0.001** | -0.20 | 0.04 | -4.84 | **<0.001** |
| Aridity | -0.00004 | 0.00004 | 1.22 | 0.225 | -0.0001 | 0.0002 | -0.63 | 0.534 |
| Intercept | 0.76 | 0.01 | 68.21 | **<0.001** | 0.18 | 0.06 | 3.18 | **0.003** |
| $R^2$ | 0.97 | | | | 0.16 | | | |

***Disturbance type***

| | **50 years (n = 138)** | | | | **100 years (n = 43)** | | | |
|---|---|---|---|---|---|---|---|---|
| **Predictor variable** | **Estimate** | **SE** | ***t*-statistic** | ***p*-value** | **Estimate** | **SE** | ***t*-statistic** | ***p*-value** |
| (LMM intercept) | -0.041 | 0.007 | -5.70 | **<0.001** | -0.208 | 0.03 | -6.03 | **<0.001** |
| Agriculture | 0.006 | 0.007 | 0.84 | 0.401 | 0.007 | 0.02 | 0.35 | 0.732 |
| Cultivation | 0.008 | 0.005 | 1.76 | 0.081 | - | - | - | - |
| Grazing | 0.017 | 0.006 | 2.64 | **0.009** | - | - | - | - |
| Intercept | 0.770 | 0.011 | 67.72 | **<0.001** | 0.189 | 0.06 | 3.22 | **0.002** |
| $R^2$ | 0.97 | | | | 0.19 | | | |

***Restoration strategy***

| | **50 years (n = 144)** | | | | **100 years (n = 46)** | | | |
|---|---|---|---|---|---|---|---|---|
| **Predictor variable** | **Estimate** | **SE** | ***t*-statistic** | ***p*-value** | **Estimate** | **SE** | ***t*-statistic** | ***p*-value** |
| (LMM intercept) | -0.042 | 0.007 | -5.87 | **<0.001** | -0.21 | 0.03 | -6.21 | **<0.001** |
| Active | -0.008 | 0.006 | -1.34 | 0.184 | 0.02 | 0.03 | -0.62 | 0.537 |
| Intercept | 0.760 | 0.012 | 65.48 | **<0.001** | 0.18 | 0.06 | 3.16 | **0.003** |
| $R^2$ | 0.97 | | | | 0.16 | | | |

***Life form***

| | **50 years (n = 128)** | | | | **100 years (n = 49)** | | | |
|---|---|---|---|---|---|---|---|---|
| **Predictor variable** | **Estimate** | **SE** | **t-statistic** | ***p*-value** | **Estimate** | **SE** | ***t*-statistic** | ***p*-value** |

| | | | | | | | | |
|---|---|---|---|---|---|---|---|---|
| (LMM intercept) | -0.047 | 0.008 | -5.95 | <**0.001** | -0.231 | 0.03 | -8.10 | <**0.001** |
| Fungi | -0.002 | 0.008 | -0.24 | 0.813 | 0.016 | 0.02 | 0.81 | 0.425 |
| Invertebrate | 0.002 | 0.004 | 0.44 | 0.659 | 0.010 | 0.02 | 0.46 | 0.646 |
| Microorganism | -0.002 | 0.011 | -0.17 | 0.865 | 0.013 | 0.02 | 0.55 | 0.583 |
| Non-woody | -0.012 | 0.009 | -1.30 | 0.195 | 0.005 | 0.02 | 0.22 | 0.831 |
| Intercept | 0.750 | 0.013 | 60.94 | <**0.001** | 0.148 | 0.05 | 3.03 | **0.004** |
| $R^2$ | 0.97 | | | | 0.16 | | | |

**Table S13.** Results of the linear models fitted for the effect of aridity, disturbance category, restoration strategy and life form on the response ratio predicted for species similarity after 50 and 100 years of recovery. The reference level for the disturbance type is "logging" and for the life form is "woody". Bold *p*-values were considered significant.

| ***Aridity*** | | | | | | | | |
|---|---|---|---|---|---|---|---|---|
| | **50 years (n = 35)** | | | | **100 years (n = 20)** | | | |
| **Predictor variable** | **Estimate** | **SE** | ***t*-statistic** | ***p*-value** | **Estimate** | **SE** | ***t*-statistic** | ***p*-value** |
| (LMM intercept) | -0.09 | 0.06 | -1.57 | 0.125 | -0.10 | 0.05 | -1.94 | 0.070 |
| Aridity | 0.0001 | 0.0003 | 0.27 | 0.789 | 0.00006 | 0.0004 | 0.16 | 0.877 |
| Intercept | 0.77 | 0.04 | 17.35 | **<0.001** | 0.61 | 0.04 | 16.97 | **<0.001** |
| $R^2$ | 0.90 | | | | 0.94 | | | |

| ***Disturbance type*** | | | | | | | | |
|---|---|---|---|---|---|---|---|---|
| | **50 years (n = 31)** | | | | **100 years (n = 16)** | | | |
| **Predictor variable** | **Estimate** | **SE** | ***t*-statistic** | ***p*-value** | **Estimate** | **SE** | ***t*-statistic** | ***p*-value** |
| (LMM intercept) | -0.10 | 0.04 | -2.69 | **0.012** | -0.096 | 0.039 | -2.48 | **0.025** |
| Agriculture | 0.02 | 0.03 | 0.56 | 0.582 | 0.004 | 0.024 | 0.18 | 0.857 |
| Cultivation | 0.02 | 0.03 | 0.58 | 0.569 | -0.017 | 0.052 | -0.32 | 0.753 |
| Grazing | 0.16 | 0.04 | 3.93 | **<0.001** | - | - | - | - |
| Intercept | 0.77 | 0.04 | 20.14 | **<0.001** | 0.614 | 0.037 | 16.44 | **<0.001** |
| $R^2$ | 0.94 | | | | 0.93 | | | |

| ***Life form*** | | | | | | | | |
|---|---|---|---|---|---|---|---|---|
| | **50 years (n = 35)** | | | | **100 years (n = 20)** | | | |
| **Predictor variable** | **Estimate** | **SE** | **t-statistic** | ***p*-value** | **Estimate** | **SE** | ***t*-statistic** | ***p*-value** |
| (LMM intercept) | -0.056 | 0.042 | -1.34 | 0.192 | -0.100 | 0.042 | -2.30 | **0.038** |
| Fungi | -0.007 | 0.073 | -0.10 | 0.921 | -0.006 | 0.055 | -0.11 | 0.911 |
| Invertebrate | -0.017 | 0.031 | -0.54 | 0.591 | -0.006 | 0.034 | 0.02 | 0.986 |
| Microorganism | -0.004 | 0.053 | -0.08 | 0.936 | -0.001 | 0.041 | 0.03 | 0.980 |
| Non-woody | -0.001 | 0.044 | -0.03 | 0.978 | 0.011 | 0.041 | 0.275 | 0.787 |
| Intercept | 0.801 | 0.041 | 19.43 | **<0.001** | 0.614 | 0.040 | 15.25 | **<0.001** |
| $R^2$ | 0.92 | | | | 0.93 | | | |

**Table S14.** Results of the linear models fitted for the effect of aridity, disturbance category and restoration strategy on the response ratio predicted for carbon cycling after 50 and 100 years of recovery. The reference level for the disturbance type is “logging” and for the restoration strategy is “passive”. Bold *p*-values were considered significant.

***Aridity***

| | 50 years (n = 122) | | | | 100 years (n = 45) | | | |
|---|---|---|---|---|---|---|---|---|
| **Predictor variable** | **Estimate** | **SE** | ***t*-statistic** | ***p*-value** | **Estimate** | **SE** | ***t*-statistic** | ***p*-value** |
| (LMM intercept) | -0.06 | 0.012 | -4.99 | **<0.001** | -0.23 | 0.065 | -3.51 | **<0.001** |
| Aridity | -0.00004 | 0.00003 | -1.14 | 0.258 | -0.0004 | 0.0003 | -1.41 | 0.166 |
| Intercept | 0.78 | 0.016 | 47.81 | **<0.001** | 0.27 | 0.087 | 3.10 | **0.003** |
| $R^2$ | 0.26 | | | | 0.26 | | | |

***Disturbance type***

| | 50 years (n = 120) | | | | 100 years (n = 43) | | | |
|---|---|---|---|---|---|---|---|---|
| **Predictor variable** | **Estimate** | **SE** | ***t*-statistic** | ***p*-value** | **Estimate** | **SE** | ***t*-statistic** | ***p*-value** |
| (LMM intercept) | -0.088 | 0.011 | -7.99 | **<0.001** | -0.38 | 0.06 | -6.30 | **<0.001** |
| Agriculture | 0.003 | 0.023 | 0.17 | 0.865 | 0.02 | 0.14 | 0.17 | 0.863 |
| Cultivation | 0.029 | 0.008 | 3.78 | **<0.001** | 0.16 | 0.04 | 3.66 | **<0.001** |
| Grazing | 0.029 | 0.011 | 2.59 | **0.011** | - | - | - | - |
| Intercept | 0.770 | 0.015 | 50.55 | **<0.001** | 0.25 | 0.07 | 3.34 | **0.002** |
| $R^2$ | 0.96 | | | | 0.41 | | | |

***Restoration strategy***

| | 50 years (n = 122) | | | | 100 years(n = 45) | | | |
|---|---|---|---|---|---|---|---|---|
| **Predictor variable** | **Estimate** | **SE** | ***t*-statistic** | ***p*-value** | **Estimate** | **SE** | ***t*-statistic** | ***p*-value** |
| (LMM intercept) | -0.067 | 0.011 | -6.11 | **<0.001** | -0.26 | 0.06 | -4.19 | **<0.001** |
| Active | -0.007 | 0.017 | -0.45 | 0.651 | 0.04 | 0.07 | 0.56 | 0.580 |
| Intercept | 0.777 | 0.016 | 47.42 | **<0.001** | 0.31 | 0.09 | 3.65 | **<0.001** |
| $R^2$ | 0.95 | | | | 0.23 | | | |

**Table S15.** Results of the linear models fitted for the effect of aridity, disturbance category and restoration strategy on the response ratio predicted for nitrogen stock after 50 and 100 years of recovery. The reference level for the disturbance type is "logging" and for the restoration strategy is "passive". Bold *p*-values were considered significant.

| ***Aridity*** | | | | | | | | |
|---|---|---|---|---|---|---|---|---|
| | **50 years (n = 76)** | | | | **100 years (n = 26)** | | | |
| **Predictor variable** | **Estimate** | **SE** | ***t*-statistic** | ***p*-value** | **Estimate** | **SE** | ***t*-statistic** | ***p*-value** |
| (LMM intercept) | -0.01 | 0.01 | -0.98 | 0.329 | -0.07 | 0.05 | -1.46 | 0.159 |
| Aridity | -0.00006 | 0.00002 | -2.57 | **0.012** | -0.0003 | 0.0001 | -0.23 | 0.823 |
| Intercept | 0.84 | 0.02 | 39.33 | **<0.001** | 0.60 | 0.08 | 7.77 | **<0.001** |
| $R^2$ | 0.96 | | | | 0.71 | | | |

| ***Disturbance type*** | | | | | | | | |
|---|---|---|---|---|---|---|---|---|
| | **50 years (n = 75)** | | | | **100 years (n = 25)** | | | |
| **Predictor variable** | **Estimate** | **SE** | ***t*-statistic** | ***p*-value** | **Estimate** | **SE** | ***t*-statistic** | ***p*-value** |
| (LMM intercept) | -0.036 | 0.012 | -2.98 | **0.004** | -0.066 | 0.047 | -1.40 | **0.177** |
| Agriculture | 0.023 | 0.012 | 1.91 | 0.060 | 0.010 | 0.028 | 0.36 | 0.721 |
| Cultivation | 0.021 | 0.007 | 3.03 | **0.003** | 0.004 | 0.021 | 0.17 | 0.869 |
| Grazing | 0.006 | 0.011 | 0.50 | 0.617 | - | - | - | - |
| Intercept | 0.839 | 0.021 | 39.25 | **<0.001** | 0.613 | 0.089 | 6.90 | **<0.001** |
| $R^2$ | 0.96 | | | | 0.69 | | | |

| ***Restoration strategy*** | | | | | | | | |
|---|---|---|---|---|---|---|---|---|
| | **50 years (n = 76)** | | | | **100 years (n = 26)** | | | |
| **Predictor variable** | **Estimate** | **SE** | ***t*-statistic** | ***p*-value** | **Estimate** | **SE** | ***t*-statistic** | ***p*-value** |
| (LMM intercept) | -0.029 | 0.012 | -2.37 | **0.021** | -0.07 | 0.04 | -1.58 | 0.128 |
| Active | 0.001 | 0.011 | 0.12 | 0.905 | -0.02 | 0.02 | -1.12 | 0.273 |
| Intercept | 0.831 | 0.022 | 38.14 | **<0.001** | 0.60 | 0.08 | 7.94 | **<0.001** |
| $R^2$ | 0.95 | | | | 0.72 | | | |

**Table S16.** Results of the linear models fitted for the effect of aridity, disturbance category and restoration strategy on the response ratio predicted for phosphorus stock after 50 years of recovery. The reference level for the disturbance type is "logging" and for the restoration strategy is "passive". Bold *p*-values were considered significant.

| ***Aridity* (n = 24)** | | | | |
|---|---|---|---|---|
| | **50 years** | | | |
| **Predictor variable** | **Estimate** | **SE** | ***t*-statistic** | ***p*-value** |
| (LMM intercept) | 0.07 | 0.01 | 5.41 | **<0.001** |
| Aridity | -0.000009 | 0.00002 | -0.38 | 0.707 |
| Intercept | 1.05 | 0.02 | 54.63 | **<0.001** |
| $R^2$ | 0.99 | | | |
| ***Disturbance type* (n = 24)** | | | | |
| | **50 years** | | | |
| **Predictor variable** | **Estimate** | **SE** | ***t*-statistic** | ***p*-value** |
| (LMM intercept) | 0.070 | 0.013 | 5.57 | **<0.001** |
| Agriculture | 0.003 | 0.020 | 0.18 | 0.862 |
| Cultivation | 0.005 | 0.010 | 0.47 | 0.643 |
| Intercept | 1.051 | 0.020 | 51.50 | **<0.001** |
| $R^2$ | 0.99 | | | |
| ***Restoration strategy* (n = 24)** | | | | |
| | **50 years** | | | |
| **Predictor variable** | **Estimate** | **SE** | ***t*-statistic** | ***p*-value** |
| (LMM intercept) | -0.029 | 0.012 | -2.37 | **0.021** |
| Active | 0.001 | 0.011 | 0.12 | 0.905 |
| Intercept | 0.831 | 0.022 | 38.14 | **<0.001** |
| $R^2$ | 0.95 | | | |

**Table S17.** Pairwise comparisons among the categories of the variable "disturbance category" when they had a significant effect on the predicted response ratio of each recovery metric after 50 and 100 years of recovery. Bold *p*-values were considered significant.

| Comparisons | Estimate | SE | *t*-ratio | *p*-value | value ± SE | CI (0.95) |
|---|---|---|---|---|---|---|
| **ORGANISM ABUNDANCE** | | | | | | |
| **50 years** | | | | | | |
| **Comparisons** | **Estimate** | **SE** | ***t*-ratio** | ***p*-value** | **value ± SE** | **CI (0.95)** |
| Logging | | | | | -0.649 ± 0.007 | [-0.66, -0.64] |
| Logging – Agriculture | -0.003 | 0.017 | -0.24 | 0.995 | | |
| Logging – Cultivation | -0.007 | 0.011 | -0.66 | 0.911 | | |
| Logging – Grazing | -0.033 | 0.013 | -2.60 | **0.049** | | |
| Agriculture | | | | | -0.646 ± 0.012 | [-0.68, -0.62] |
| Agriculture – Cultivation | -0.004 | 0.014 | -0.27 | 0.993 | | |
| Agriculture – Grazing | -0.030 | 0.016 | -1.91 | 0.228 | | |
| Cultivation | | | | | -0.642 ± 0.008 | [-0.66, -0.63] |
| Cultivation – Grazing | -0.026 | 0.013 | -2.03 | 0.183 | | |
| Grazing | | | | | -0.616 ± 0.010 | [-0.64, -0.60] |
| **SPECIES DIVERSITY** | | | | | | |
| **50 years** | | | | | | |
| **Comparisons** | **Estimate** | **SE** | ***t*-ratio** | ***p*-value** | **value ± SE** | **CI (0.95)** |
| Logging | | | | | -0.511 ± 0.003 | [-0.52, -0.51] |
| Logging – Agriculture | -0.006 | 0.016 | -2.13 | 0.835 | | |
| Logging – Cultivation | -0.008 | 0.004 | -1.34 | 0.297 | | |
| Logging – Grazing | -0.017 | 0.006 | -2.53 | **0.045** | | |
| Agriculture | | | | | -0.505 ± 0.007 | [-0.52, -0.49] |
| Agriculture – Cultivation | -0.002 | 0.017 | 1.75 | 0.992 | | |
| Agriculture – Grazing | -0.011 | 0.009 | -1.22 | 0.614 | | |
| Cultivation | | | | | -0.503 ± 0.004 | [-0.51, -0.50] |
| Cultivation – Grazing | -0.009 | 0.007 | -1.27 | 0.587 | | |
| Grazing | | | | | -0.494 ± 0.006 | [-0.51, -0.48] |

| SIMILARITY | | | | | | |
|---|---|---|---|---|---|---|
| **50 years** | | | | | | |
| **Comparisons** | **Estimate** | **SE** | ***t*-ratio** | ***p*-value** | **value ± SE** | **CI (0.95)** |
| Logging | | | | | -0.800 ± 0.015 | [-0.83, -0.77] |
| Logging – Agriculture | -0.016 | 0.029 | -0.56 | 0.944 | | |
| Logging – Cultivation | -0.019 | 0.032 | -0.58 | 0.939 | | |
| Logging – Grazing | -0.159 | 0.041 | -3.93 | **0.003** | | |
| Agriculture | | | | | -0.784 ± 0.025 | [-0.83, -0.73] |
| Agriculture – Cultivation | -0.003 | 0.037 | -0.07 | 1.000 | | |
| Agriculture – Grazing | -0.143 | 0.045 | -3.19 | **0.017** | | |
| Cultivation | | | | | -0.781 ± 0.028 | [-0.84, -0.72] |
| Cultivation – Grazing | -0.141 | 0.047 | -3.00 | **0.026** | | |
| Grazing | | | | | -0.641 ± 0.038 | [-0.72, -0.56] |

| CARBON CYCLING | | | | | | |
|---|---|---|---|---|---|---|
| **50 years** | | | | | | |
| **Comparisons** | **Estimate** | **SE** | **t-ratio** | **p-value** | **value ± SE** | **CI (0.95)** |
| Logging | | | | | -0.570 ± 0.006 | [-0.58, -0.56] |
| Logging – Agriculture | -0.003 | 0.023 | -0.17 | 1.000 | | |
| Logging – Cultivation | -0.029 | 0.008 | -3.78 | **0.001** | | |
| Logging – Grazing | -0.029 | 0.011 | -2.59 | **0.052** | | |
| Agriculture | | | | | -0.566 ± 0.022 | [-0.61, -0.52] |
| Agriculture – Cultivation | -0.026 | 0.022 | -1.14 | 0.666 | | |
| Agriculture – Grazing | -0.025 | 0.024 | -1.06 | 0.718 | | |
| Cultivation | | | | | -0.541 ± 0.005 | [-0.55, -0.53] |
| Cultivation – Grazing | 0.0004 | 0.011 | 0.04 | 1.000 | | |
| Grazing | | | | | -0.541 ± 0.009 | [-0.56, -0.52] |
| **100 years** | | | | | | |
| Logging | | | | | -0.541 ± 0.033 | [-0.61, -0.47] |
| Logging – Agriculture | -0.024 | 0.140 | -0.17 | 0.983 | | |

| | | | | | | |
|---|---|---|---|---|---|---|
| Logging – Cultivation | -0.160 | 0.043 | -3.66 | **0.002** | | |
| Agriculture | | | | | -0.541 ± 0.033 | [-0.79, -0.24] |
| Agriculture – Cultivation | -0.135 | 0.138 | -0.98 | 0.595 | | |
| Cultivation | | | | | -0.541 ± 0.033 | [-0.44, -0.32] |
| **NITROGEN STOCK** | | | | | | |
| **50 years** | | | | | | |
| **Comparisons** | **Estimate** | **SE** | **t-ratio** | **p-value** | **value ± SE** | **CI (0.95)** |
| Logging | | | | | -0.488 ± 0.005 | [-0.50, -0.48] |
| Logging – Agriculture | -0.023 | 0.012 | -1.91 | 0.233 | | |
| Logging – Cultivation | -0.209 | 0.007 | -3.03 | **0.018** | | |
| Logging – Grazing | -0.006 | 0.011 | -0.50 | 0.958 | | |
| Agriculture | | | | | -0.465 ± 0.011 | [-0.49, -0.44] |
| Agriculture – Cultivation | 0.002 | 0.012 | 0.20 | 0.997 | | |
| Agriculture – Grazing | 0.018 | | 1.16 | 0.652 | | |
| Cultivation | | | | | -0.467 ± 0.005 | [-0.48, -0.46] |
| Cultivation – Grazing | 0.015 | 0.015 | 1.36 | 0.530 | | |
| Grazing | | 0.011 | | | -0.482 ± 0.010 | [-0.50, -0.46] |
